# PSII photoinhibition as a protective strategy against PSI photoinhibition: Maintaining PSI in an oxidized state by suppressing PSII activity under environmental stress

**DOI:** 10.1101/2025.05.06.652500

**Authors:** Ko Takeuchi, Shintaro Harimoto, Shu Maekawa, Chikahiro Miyake, Kentaro Ifuku

## Abstract

Photosystem I (PSI) can be photoinhibited by excessive electron flow from Photosystem II (PSII), causing serious growth inhibition due to PSI’s limited repair capacity. In contrast, PSII is more susceptible to photoinhibition under stress but has an efficient repair system. Consequently, PSII photoinhibition is considered a protective mechanism that mitigates PSI over-reduction. However, this photoprotective role under environmental stress remains unexplored in intact plants without using mutants or chemical treatments. To address this, we examined the relationship between PSII photoinhibition and susceptibility to PSI photoinhibition under two representative stresses that selectively induce PSI photoinhibition: chilling stress and fluctuating light, using *A. thaliana* and cucumber. Under chilling stress, *A. thaliana* exhibited marked PSII photoinhibition and maintained active PSI, whereas cucumber showed insufficient PSII downregulation and suffered PSI photoinhibition. In addition, when fluctuating light treatment was applied to plants with various Fv/Fm (the maximum quantum yield of PSII), plants with reduced Fv/Fm maintained oxidized PSI, and PSI photoinhibition progressed slowly. The susceptibility to PSI photoinhibition under fluctuating light strongly correlated with Fv/Fm, providing clear evidence that PSII photoinhibition protects PSI *in vivo*. Interestingly, even in plants where P700 remained oxidized during saturation pulses due to PSII photoinhibition, the iron-sulfur clusters (Fe-S clusters) remained reduced. However, the re-oxidation rate of reduced Fe-S clusters was enhanced in PSII-photoinhibited plants, suggesting that charge recombination to P700^+^ with reduced components on the acceptor side would suppress ROS generation downstream of PSI. This study clarifies how PSII photoinhibition suppresses PSI photoinhibition and prevents ROS-induced damage in wild-type plants under environmental stress.

## 1 Introduction

Plants are exposed to diverse environmental stresses in nature. Many environmental stresses, such as temperature, drought, and salinity stress, cause photosystem II (PSII) photoinhibition (Powles, 1984). Some stresses such as fluctuating light (Tsuyama and Kobayashi, 2009; Terashima et al., 2021; Basso et al., 2022; Gollan et al., 2023; Sun et al., 2023; Yang et al., 2024) and low-temperature stress in cucumber (Terashima et al., 1994; Nakano et al., 2010; Sonoike, 2011; Takeuchi et al., 2022, 2024) and various other plants (Havaux and Davaud, 1994; Ivanov et al., 1998; Kim et al., 2005; Ivanov et al., 2012a, 2012b; Chen et al., 2024; Li et al., 2024) induce photosystem I (PSI) photoinhibition. The turnover of certain proteins in PSII is considerably faster than that in PSI (Melis, 1999; Miyata et al., 2012; Li et al., 2018; Lima-Melo et al., 2019), indicating that PSII photoinhibition is prioritized over PSI photoinhibition to mitigate the adverse effects on plant growth under diverse stress conditions (Barber and Andersson, 1992; Suzuki et al., 2021).

The main actor in photoinhibition is reactive oxygen species (ROS) (Asada, 2006; Rutherford et al., 2012; Degen and Johnson, 2024). On one hand, ROS act as important secondary messengers that trigger plant acclimation responses via gene expression, but uncontrollable levels of ROS in the photosystems induce photoinhibition (Takahashi and Asada, 1988; Asada, 2006; Murata et al., 2007; Sadiq et al., 2019; Dorion et al., 2021; Hamada et al., 2023; Wang et al., 2024). In PSII, the core protein is damaged by singlet oxygen (^1^O_2_) originating from an excited triplet state of P_680_ (^3^P_680_*) and charge recombination reactions between the secondary quinone acceptor, Q_B_, and the S_2,3_ state of the Mn cluster (Krieger-Liszkay, 2005; Tyystjärvi, 2008; Vass, 2011; Rutherford et al., 2012; Kumar et al., 2021). Additionally, PSII reduces oxygen to generate O_2_^•–^ and H_2_O_2_ both on the PSII electron donor and acceptor sides (Pheo, Q_A_, Q_B_, plastoquinone, and Cyt *b_559_*) (Kale et al., 2017). H_2_O_2_ is converted to hydroxyl radicals (OH^•^) in the presence of free Fe or around non-heme Fe in PSII, which subsequently generate lipid peroxide radicals (LOO^•^), contributing to PSII photoinhibition (Pospíšil and Yamamoto, 2017; Kumar et al., 2021; Foyer and Hanke, 2022). Furthermore, the PSII recovery system is susceptible to ROS (Nishiyama et al., 2001, 2006; Murata et al., 2007). In PSI, the redox potential of electron carriers is sufficiently low to reduce O_2_ to O_2_^•–^, allowing direct electron transfer to O_2_ (Mehler, 1951; Cherepanov et al., 2017; Hani et al., 2024). Furthermore, the dismutation of O_2_^•–^ into H_2_O_2_ leads to the generation of OH^•^ via the Fenton reaction around iron-sulfur clusters (Fe-S clusters) (Petronek et al., 2021). Additionally, when the acceptor side of PSI is fully reduced, recombination between radical pairs P700^+^/A_0_^−^ or P700^+^/A_1_^−^ can generate a triplet state of P700, which produces ^1^O_2_ (Shuvalov et al., 1986; Rutherford et al., 2012; Degen and Johnson, 2024). These ROS lead to a decrease in photo-oxidizable P700 (PSI photoinhibition) due to destruction of Fe-S clusters and damage to the PSI core (Inoue et al., 1989; Sonoike and Terashima, 1994; Sonoike et al., 1995; Sonoike, 1996; Erling Tjus et al., 1998; Nakano et al., 2006; Kılıç et al., 2023; Shimakawa et al., 2024).

However, the production site of ROS that induces PSI photoinhibition is still under discussion (Takahashi and Asada, 1988; Kozuleva et al., 2014; Kozuleva et al., 2021; Shimakawa et al., 2024).

To protect PSI from photoinhibition, the accumulation of oxidized PSI reaction center chlorophyll P700 (P700^+^) is important (Trissl, 1997; Shimakawa and Miyake, 2018a; Miyake, 2020). Plants have various mechanisms to maintain P700 in an oxidized state under environmental stress conditions. On the electron acceptor side, the proper activity of the Calvin-Benson-Bassham (CBB) cycle and photorespiration in C_3_ plants serve as crucial electron sinks for oxidizing P700 (Hanawa et al., 2017; Furutani et al., 2020; Wada et al., 2020). Photosynthetic organisms, except for angiosperms, have flavodiiron proteins (FLV) that mediate electron flow from PSI to oxygen, contributing to P700 oxidation (Helman et al., 2003; Helman et al., 2005; Shimakawa et al., 2019; Storti et al., 2020; Bag et al., 2023). The water-water cycle also contributes to P700 oxidation (Miyake, 2010; Huang et al., 2019). On the electron donor side of PSI, acidification of the thylakoid lumen (ΔpH) suppresses electron transport in Cyt *b*_6_*f*, promoting P700 oxidation (Foyer et al., 1990; Anderson, 1992; Nishio and Whitmarsh, 1993; Hope, 2000; Baker et al., 2007; Schöttler and Tóth, 2014; Schöttler et al., 2015). ΔpH is primarily regulated by the activity of cyclic electron flow (CEF) and chloroplast ATPase (Heber and Walker, 1992; Shikanai, 2007; Kohzuma et al., 2009; Miyake, 2010; Ifuku et al., 2011; Kanazawa et al., 2017; Takagi et al., 2017a; Shikanai, 2023). Over-reduction of the plastoquinone pool (PQ) inhibits the Q-cycle of the Cyt *b*_6_*f* and contributes to the P700 oxidation, a mechanism known as the reduction-induced suppression of electron transport (RISE) (Shaku et al. 2016). The plastid terminal oxidase PTOX also protects PSI by inducing P700^+^ (Messant et al., 2024).

In addition to these P700 oxidation systems, PSII photoinhibition has been proposed to maintain P700 in an oxidized state (e.g., Ivanov et al., 1998; Kim et al., 2005). In recent years, a number of studies have shown that PSII photoinhibition promotes PSI oxidation, greatly advancing our understanding of PSI photoinhibition (Suorsa et al., 2012; Tikkanen et al., 2014; Zhang et al., 2014; Zhang et al., 2016; Obara et al., 2022; Messant et al., 2024; Napaumpaiporn et al., 2024; Ozawa et al., 2024). However, studies primarily aimed to elucidate the role of PSII photoinhibition in P700 oxidation have mainly relied on chemical treatments and mutants, such as DCMU, lincomycin, and *pgr5* mutants. A key limitation of these approaches is the uncertainty regarding whether they truly reflect the physiological and strategic responses of wild-type (WT) plants. For instance, *pgr5* mutants exhibit PSI photoinhibition even under moderate light, making them extreme models that might not represent typical stress responses in WT plants. Similarly, artificial suppression of PSII activity via chemical treatments induces PSI oxidation, but this does not necessarily mimic naturally occurring PSII photoinhibition under environmental stress.

Against this background, the present study aimed to investigate how PSII photoinhibition contributes to PSI protection under environmental stress in WT plants. To this end, we focused on two representative environmental conditions known to induce PSI photoinhibition in nature: chilling stress and fluctuating light stress. In experiments under chilling stress, we compared *A. thaliana*, a chilling-tolerant species that does not exhibit PSI photoinhibition under low temperatures at moderate light intensity, with cucumber, a chilling-sensitive species that undergoes PSI photoinhibition. Furthermore, to assess the protective effect of PSII photoinhibition on PSI, we exposed plants with prior PSII photoinhibition to fluctuating light stress. These approaches allow us to evaluate whether PSII photoinhibition mitigates PSI photoinhibition under environmental stress.

## 2 Materials and Methods

### 2.1 Plant materials, growth conditions, and light-chilling stress treatments

*A. thaliana* wild type (ecotype Columbia-0) was grown in soil under long-day conditions (16-h light/8- h dark) at 22 °C with a light intensity of 80–100 μmol photons m^-2^ s^-1^. Rosette leaves from 4- to 6- week-old plants were used for measurements. Cucumber plants (cv High Green 21, Saitama Genshu Ikuseikai, Japan) were grown in plastic pots with soil under long-day conditions (16-h light/8-h dark) at 27 °C in a growth chamber. Fully expanded leaves from plants 3 to 4 weeks after germination were used for analyses.

Four- to six-week-old *A. thaliana* plants were subjected to chilling stress at 4 °C in a cold room under light intensities of 180–250 µmol photons m^-2^ s^-1^ for 1–4 h (short-term chilling stress) or more than 60 h (long-term chilling stress). After chilling treatment, plants were dark-adapted at 4 °C for at least 20 min. Subsequently, photosynthetic parameters were measured at room temperature (23 °C or 25 °C) within 5 min of transferring the plants from the cold room. Plants without chilling treatment were used as the controls. Leaf discs prepared from fully expanded leaves of 3- to 4-week-old cucumber plants were floated on tap water (adaxial side facing upward) and exposed to chilling stress at 4 °C in a cold room under light intensities ranging from 80–650 µmol photons m^-2^ s^-1^ for 20 h.

### 2.2 Measurement of chlorophyll fluorescence, P700, iron/sulfur clusters signals, and gas-exchange

Chlorophyll fluorescence and redox change of P700 were measured using induction curve methods with Dual-PAM 100 (Heinz Walz GmbH, Effeltrich, Germany) at room temperature as previously described (Takeuchi et al., 2022; Takeuchi et al., 2024). The maximum quantum yield of PSII in the dark was calculated as Fv/Fm. The effective quantum yield of PSII was calculated as Y(II) = (Fm’− Fs)/Fm’. The quantum yield of non-regulated energy dissipation in PSII was calculated as Y(NO) = Fs/Fm. The quantum yield of regulated energy dissipation in PSII was calculated as Y(NPQ) = Fs/Fm’ − Y(NO). The fraction of open PSII reaction centers in a lake model was calculated as qL = (Fm’− Fs)/(Fm’– Fó) × Fó/Fs. The Fo and Fm levels, the minimum and maximum fluorescence yield, respectively, were measured after at least 20 min of dark adaptation. The Fó (minimum fluorescence level of the illuminated sample) was calculated according to Oxborough and Baker 1997: Fó= 1/(1/Fo – 1/Fm + 1/Fm’). The light-chilling plants were adapted to darkness for at least 20 min in a cold room (4 °C), whereas the control plants were adapted to room temperature. After the onset of actinic light illumination (172 µmol photons m^-2^ s^-1^), a saturation pulse (SP, 300 ms and 20000 µmol photons m^-2^ s^-1^) was applied every 30 seconds for 5 min to monitor the Fm’ level (the maximum fluorescence yield under light illumination) and the Fs level (the steady-state fluorescence yield under light illumination).

The redox change of P700 was analyzed by monitoring the absorbance changes of transmitted light at 830 and 875 nm (Klughammer and Schreiber, 2008). The quantum yield of PSI was calculated as Y(I) = (Pm’− P)/Pm. The fraction of P700 that could not be oxidized by an SP relative to the overall P700 was calculated as Y(NA) = (Pm − Pm’)/Pm. The P700 oxidation ratio under the given actinic light was calculated as Y(ND) = P/Pm (Klughammer and Schreiber, 1994). Pm was determined by the application of an SP after the illumination with far-red (FR) light. Pm’ represents the maximum level of P700^+^ during actinic light. P represents the steady-state P700^+^ level and was recorded immediately before the application of an SP.

Rapid decay kinetics of P700^+^ under actinic light were measured using dark-interval relaxation kinetics (DIRK) analysis (Sacksteder and Kramer, 2000) to evaluate electron transfer activity from PSII to PSI. Plants before and after chilling stress were dark-adapted and illuminated with actinic light (172 µmol photons m^-2^ s^-1^) for 60 s. The kinetics of the P700^+^ were monitored after the actinic light was turned off. The levels of P700^+^ were normalized by the maximum amount of P700 (Pm) in the individual leaves.

The redox state of iron/sulfur (Fe-S) clusters, including F_X_, F_A_/F_B_, and Fd, was measured using Dual/KLAS-NIR (Heinz Walz GmbH, Effeltrich, Germany) as previously described (Klughammer and Schreiber, 2016).

CO_2_/H_2_O-exchange during the induction phase of photosynthesis was measured for 4–5 min after the onset of red actinic light (172 μmol photons m^−2^ s^−1^) at 25 °C using infra-red gas analyzer (IRGA) LI-7000 (Li-COR, Lincoln, NE, USA) measuring systems equipped with 3010-DUAL gas exchange chamber (Heinz Walz GmbH), under 40 Pa CO_2_ / 21 kPa O_2_ as described (Ohnishi et al., 2023; Maekawa et al., 2024). The gases were saturated with water vapor at 14 ± 0.1 °C. Leaf temperature was controlled at 25 ± 0.5 °C, and relative humidity was maintained at 50–60%.

### 2.3 Analysis of electron influx into P700^+^ using saturation pulses

The kinetics of P700^+^ during SP illumination after dark adaptation were obtained in the same manner as for Pm determination; after the determination of Fv/Fm, leaves before and after chilling stress were irradiated with FR for 10 s and then with an SP of 20000 µmol photons m^-2^ s^-1^ for 300 ms under FR illumination. The obtained kinetics of P700^+^ were adjusted to a maximum value of 1.

### 2.4 PSI photoinhibition (rSP) treatment

Fluctuating light treatment was applied to intact leaves (before or after chilling stress) using repetitive saturation pulse (rSP) treatment as described previously (Sejima et al., 2014; Takagi et al., 2017b). Leaves were illuminated with short pulses (20000 µmol photons m^-2^ s^-1^, 300 ms) every 10 s for 30 min in the dark (180 times SP in total). During rSP treatment, leaf temperature was maintained at 23 °C using a temperature control unit (GFS-3000; Heinz Walz GmbH, Effeltrich, Germany). After rSP treatment, the leaves were kept in darkness for 30 min, and the residual activity of PSI and PSII was calculated by measuring Pm and Fv/Fm and normalizing the values to those measured before rSP treatment as 1. For the measurement of Fv/Fm before rSP treatment, plants were dark-adapted under low-temperature (4 °C) conditions, in which the PSII repair cycle is known to be inactive (Schnettger et al., 1994).

### 2.5 Measurement of O_2_^•–^

The O_2_^•–^ content before and after rSP treatment was detected as previously described (Zhang et al., 2010). Fresh leaves were harvested before and right after rSP treatment. The liquid nitrogen-frozen samples were homogenized and mixed well with 1.5 mL phosphate buffer (pH 7.8) containing 10 μM hydroxylammonium chloride and 100 μM EDTA-Na_2_. The homogenate was centrifuged at 5000 rpm for 5 min at 4 °C and 500 μL of supernatants were transferred into fresh tubes, to which 1 ml of 17 mM sulphanilamide (in 30 % acetic acid) and 1 ml of 7 mM naphthalene diamine dihydrochloride were added in sequence, and incubated for 10 min at 37 °C. After adding 3 ml of diethyl ether into each tube, the mixtures were centrifuged at 5000 rpm for 5 min at room temperature. Absorbance at 540 nm for each sample was recorded. Calibration curves were established from 0 to 4 μM NO_2_^−^. The *r*^2^ between O.D. = 540 and [NO_2_^−^] was 0.999. From the following reaction: 2O_2_^•–^ + H^+^ + NH_2_OH → H_2_O_2_ + H_2_O + NO_2_^−^, the concentration of O_2_^•–^ was calculated from the calibration curve according to [O_2_^•–^] = 2[NO_2_^−^] (μM).

### 2.6 Statistical analysis

Statistical analyses of the data were based on Tukey-Kramer’s multiple comparison test following the one-way ANOVA or Student’s *t*-test. All calculations were performed with at least three independent biological replicates.

## 3 Results

### 3.1 Electron transfer activity and photoinhibition under chilling stress in *A. thaliana*

To investigate the progression of photoinhibition under low-temperature stress in *A. thaliana*, two distinct chilling stress treatments were applied: a short-term chilling stress of less than 4 hours and a long-term chilling stress of over 60 hours. Chilling-treated plants were dark-adapted at 4 °C, and their electron transfer activity was subsequently measured at 23 °C (Fig. 1). As for PSII parameters, Y(II), representing the effective quantum yield of PSII under light illumination, unexpectedly increased during the induction phase of photosynthesis by short-chilling stress and decreased significantly by long-chilling stress compared to untreated control plants. Y(NPQ) was reduced by both short- and long-chilling stress. Y(NO) increased dramatically after long-chilling stress. Fv/Fm decreased slightly at short-chilling stress and was significantly reduced to approximately 0.2 after long-chilling stress. 1–qL decreased after short-chilling stress, consistent with the regulated small decrease in Fv/Fm to protect PSII from photoinhibition as previously reported (Miyake et al., 2009), and increased after long-chilling stress. The low Fv/Fm and Y(II) values after long-chilling stress were attributed to extremely low Fm (Fm’) values during SP illumination, suggesting that stable charge separation in PSII was impaired due to severe PSII photo-damage by long-chilling stress.

**Figure 1.**
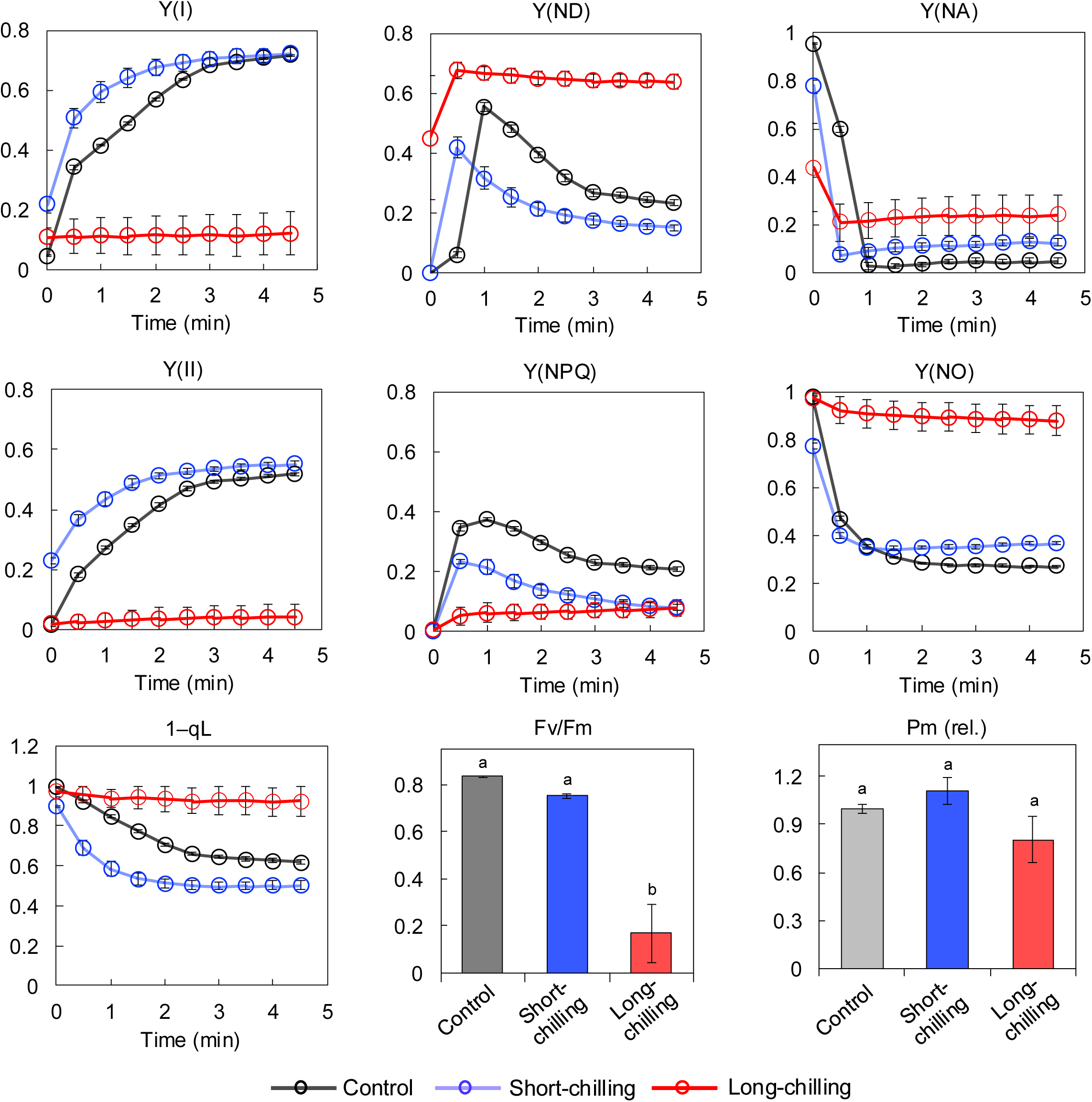
Effects of short- and long-chilling stress on photosynthetic activity in *A. thaliana*. The induction phases of chlorophyll fluorescence and chlorophyll absorption were analyzed using Dual-PAM 100. *A. thaliana* WT plants were grown for 4–6 weeks, and intact rosette leaves were analyzed before chilling stress (Control), after 4 h of chilling stress (Short-chilling), and after 96 h of chilling stress (Long-chilling). After at least 20 min of dark adaptation, the leaves were illuminated with actinic light of 172 μmol photons m^−2^ s^−1^ for 5 min under ambient conditions (23 °C) to measure Y(I), Y(ND)–the non-photochemical loss due to oxidized primary donor representing the ratio of P700^+^, Y(NA)–the non-photochemical loss due to reduced acceptor representing the acceptor side limitation of electron flow in PSI, Y(II)–the effective quantum yield of PSII under light illumination, Y(NPQ)–the yield of non-photochemical quenching, Y(NO)–the quantum yield of nonregulated energy dissipation in PSII, and qL–the fraction of open PSII reaction centers in a lake model. Fv/Fm (the maximum quantum yield of PSII) and Pm (the total amount of PSI reaction center chlorophyll P700) were calculated before the actinic light was turned on. The Pm values measured before chilling treatment were normalized to 1. Values are the mean ± SE, n = 3, biological replicates. Different letters indicate statistically significant differences by Tukey-Kramer’s multiple comparison tests after one- way ANOVA (*p* < 0.01).

As for PSI parameters, Y(ND), representing the ratio of the oxidized state of the PSI reaction center chlorophyll P700 (P700^+^), decreased slightly after short-chilling stress and increased significantly following long-chilling stress (Fig. 1). The changes in Y(ND) were due to the changes in PSII activity, as Y(II) and Y(ND) were negatively correlated (Takeuchi et al., 2022). Pm, which indicates the total amount of active PSI reaction center, showed little or no changes after short- and long-chilling stress compared to dramatic changes in Fv/Fm. These results indicated that PSII photoinhibition was induced by long-chilling stress, whereas PSI was highly oxidized, and PSI photoinhibition did not occur even after long-chilling stress.

To investigate how changes in PSII activity during chilling stress affect PSI, we adopted a novel method to evaluate electron influx from PSII to PSI by monitoring P700^+^ absorption changes during an SP immediately after dark adaptation. In the analysis of photosynthetic activity using pulse amplitude modulation (PAM), SP is applied after dark adaptation to measure the maximum level of chlorophyll fluorescence yield and the maximum P700 absorption level. During SP illumination, P700 is almost completely oxidized to P700^+^ within 5–50 ms, and subsequently reduced due to electron influx from PSII-plastocyanin (Klughammer and Schreiber, 2008a). Therefore, the reduction curve of P700^+^ during SP reflects the rate of electron donation from PSII. After short-chilling stress, where Y(II) increased during the induction phase of photosynthesis (Fig. 1), the P700^+^ decay kinetics during SP (0–300 ms) were accelerated compared to control plants (Fig. 2A), indicating enhanced electron influx into PSI. To investigate the factors responsible for the temporary increase in Y(II) (Fig. 1) and electron influx into P700^+^ after short-chilling stress (Fig. 2A), we analyzed the oxidation-reduction dynamics of Fe–S clusters during SP following dark adaptation using Dual/KLAS-NIR. Fe–S clusters were rapidly reduced upon SP illumination in both control plants and those exposed to short-chilling stress, and they remained predominantly in the reduced form throughout the 300 ms of SP as previously reported (Supplementary Fig. S1A) (Schreiber, 2017; Furutani et al., 2022). Notably, during the subsequent dark period (post-300 ms), Fe–S clusters in short-chilled plants re-oxidized more rapidly than in control plants (Supplementary Fig. S1A). The re-oxidation rate of reduced Fe–S clusters after light illumination reflects the activation level of the electron efflux reactions from Fe–S clusters (Schreiber, 2017; Kadota et al., 2019). Thus, short-term chilling stress would activate electron transfer reactions downstream of ferredoxin (Fd), thereby stimulating the entire electron transport chain, as indicated by enhanced Y(II) (Fig. 1). On the other hand, the CO₂ fixation rate during the induction phase of photosynthesis, following a few minutes of light exposure after dark adaptation, exhibited similar CO_2_ fixation ability to that in control plants (Supplementary Fig. S1B–C). These results indicated that the enhanced consumption of reducing power (Supplementary Fig. S1A) and the temporary increase in Y(II) were likely due to the activation of alternative electron flow (AEF) downstream of Fd, rather than the CBB cycle, to prevent over-reduction of the electron transport chain during the initial phase of chilling stress.

**Figure 2.**
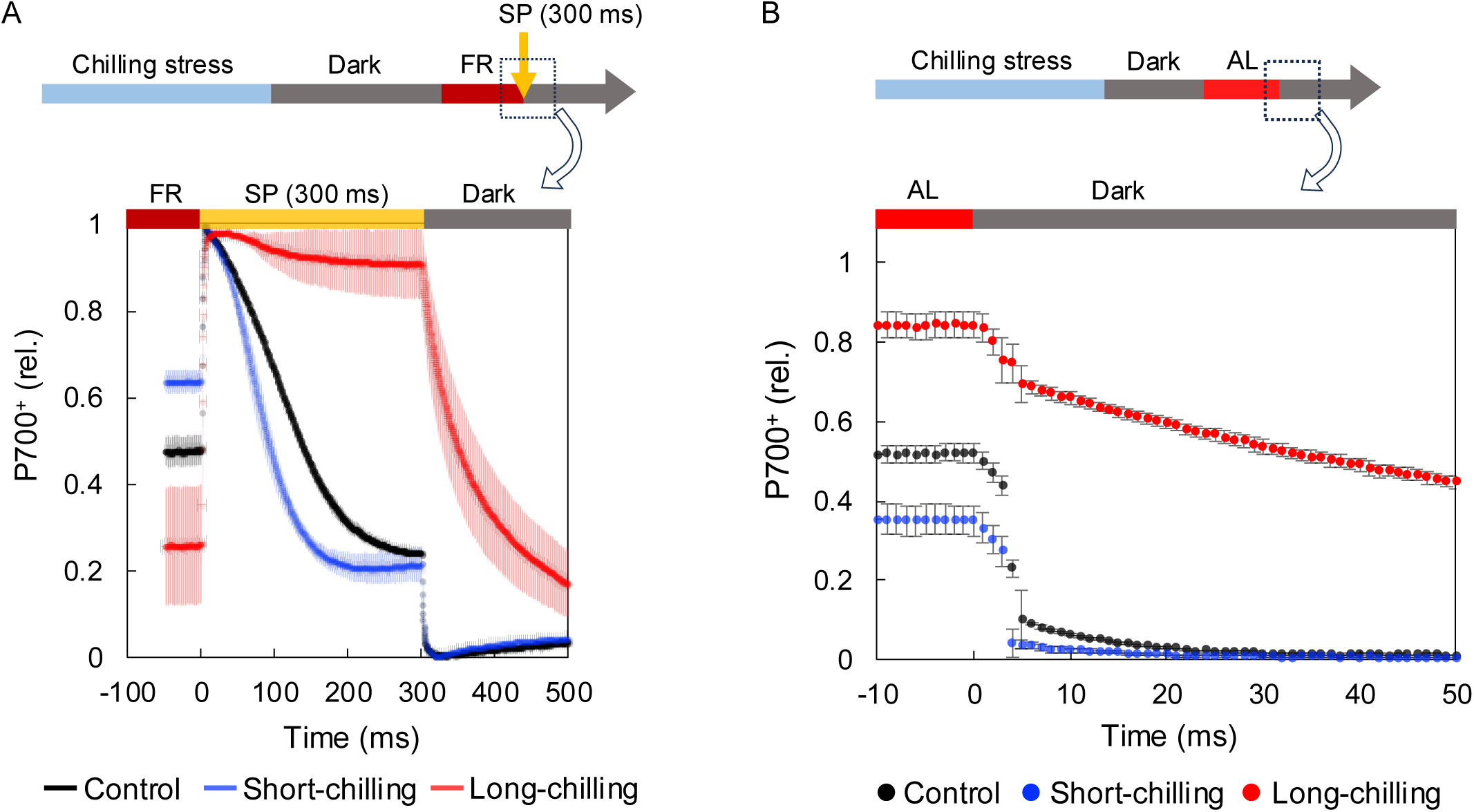
Analysis of electron influx from PSII into PSI after chilling stress in *A. thaliana*. The plants used are the same as those in Fig. 1. (A) The kinetics of P700^+^ during SP illumination were measured following at least 20 minutes of dark adaptation, using the same method as for Pm determination. Dark-adapted leaves were first irradiated with FR for 10 s and then exposed to SP illumination of 20000 µmol photons m^-2^ s^-1^ for 300 ms under ambient conditions (23 °C). The obtained P700^+^ kinetics were normalized to a maximum value of 1. Values are the mean ± SE, n = 3, biological replicates. (B) DIRK analysis of the decay kinetics of P700^+^ after actinic light was turned off. Dark- adapted chilled or non-chilled plants were illuminated with actinic light of 172 μmol photons m^−2^ s^−1^ for 1 min under ambient conditions (23 °C). After the actinic light was turned off, the decay kinetics of P700^+^ in the dark were monitored for 50 ms. The levels of P700^+^ were normalized by the maximum amount of P700 (Pm) in the individual leaves. The decay of P700^+^ is fundamentally a parameter reflecting Y(II), but care should be taken as charge-recombination fluxes could contribute to the fast component and AEF to the slow component. Values are the mean ± SE, n = 3–5, biological replicates.

Conversely, after long-chilling stress, a significant decline in PSII activity was observed (Fig. 1), leading to a marked suppression of electron influx into PSI, as indicated by the slow decay of P700⁺ during SP (Fig. 2A). DCMU treatment confirmed that the slow decay of P700⁺ during SP was due to limited electron transport from PSII (Supplementary Fig. S2) (Shimakawa et al., 2013). The suppression of electron influx into PSI by PSII photoinhibition was also validated by dark-interval relaxation kinetics (DIRK) analysis, which evaluates electron transfer activity from PSII to PSI using P700⁺ decay kinetics after turning off actinic light. After long-chilling stress, the decay of P700⁺ was significantly delayed (Fig. 2B). These results support the idea that PSII photoinhibition alleviates over-reduction of PSI and prevents PSI photoinhibition under chilling stress at moderate light intensity in *A. thaliana* (Ivanov et al., 1998; Kim et al., 2005).

### 3.2 PSI photoinhibition under chilling stress in cucumber

We conducted similar experiments using cucumber, a species known to undergo PSI photoinhibition under low-temperature conditions, unlike *A. thaliana* (Terashima et al., 1994; Kono et al., 2022). We monitored the progression of PSII and PSI photoinhibition over 24 h of chilling stress. Up to 1.5 h of chilling stress, changes in Fv/Fm and Pm were slight (Fig. 3). Importantly, after 5 h of chilling stress, Fv/Fm remained around 0.6, whereas Pm had already dropped to approximately 0.4. After 9 h of chilling stress, Fv/Fm decreased to around 0.4, whereas Pm declined to nearly 0.1. That is, the decline in Pm occurred earlier and more severely than the decline in Fv/Fm, unlike *A. thaliana*. Detailed data on electron transfer activity at each time point are shown in Supplementary Fig. S3. After 5 h of chilling stress, Y(ND) decreased, and Y(NA) increased, indicating that PSI became highly over-reduced (Supplementary Fig. S3). At the same time, the re-oxidation rate of Fe–S clusters was markedly slower in chilled cucumber compared to controls (Supplementary Fig. S4). This slower Fe- S re-oxidation rate indicates suppressed electron efflux downstream of Fd and Fe-S over-reduction (Schreiber, 2017; Takeuchi et al., in preparation). These results indicate that PSII photoinhibition by chilling stress in cucumber was not as effective as in *A. thaliana*. Consequently, PSI became over-reduced, leading to severe PSI photoinhibition. The cause of over-reduction of PSI is known to be associated with the uncoupling of thylakoid membranes at low temperatures in cucumber, leading to insufficient PSII downregulation (Peeler and Naylor, 1988; Terashima et al., 1991a, 1991b).

**Figure 3.**
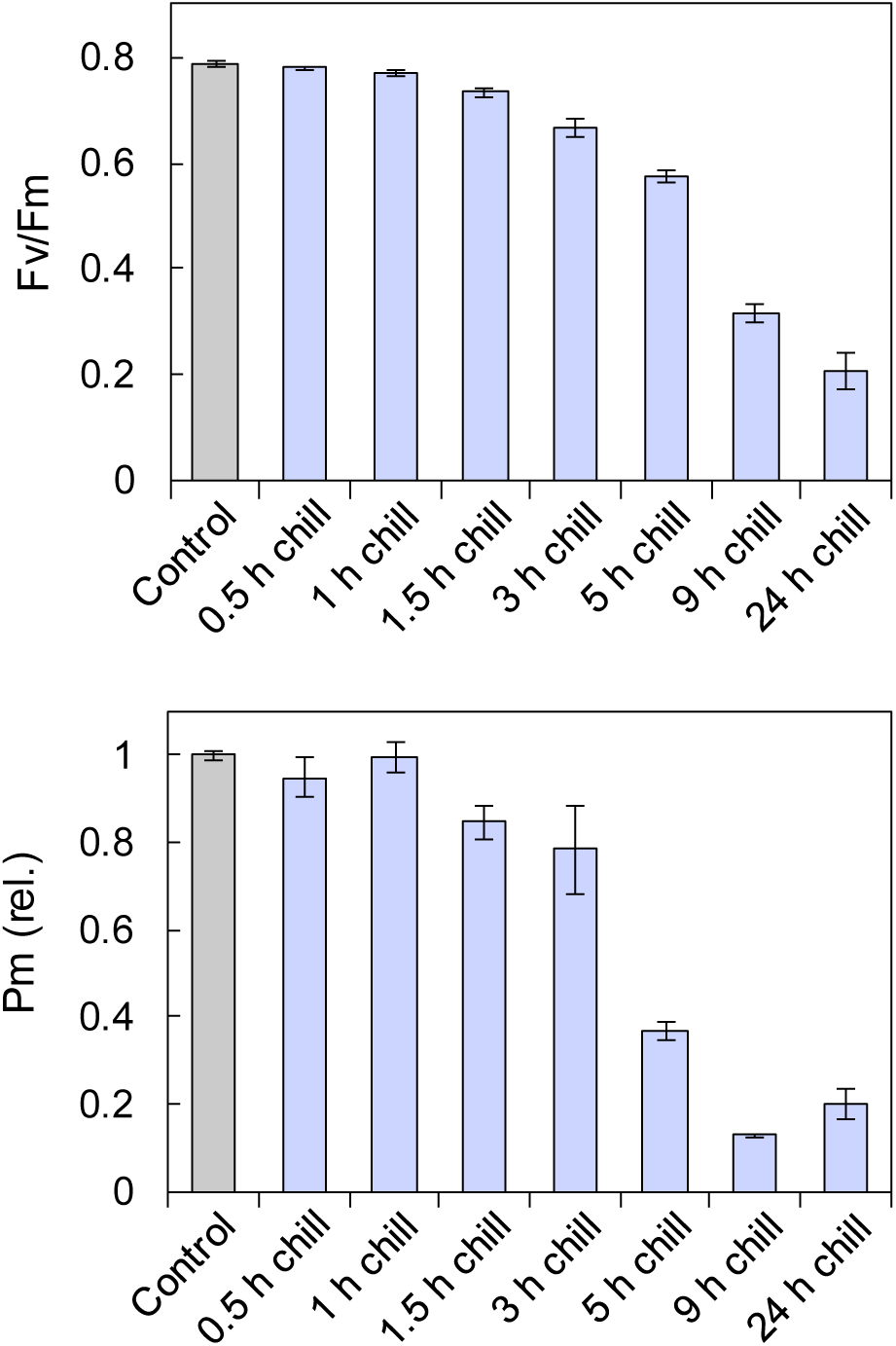
Effect of chilling stress on PSII and PSI in cucumber. Fv/Fm and Pm were analyzed in 2–3-week-old cucumber plants before (Control) and after 0.5 h–24 h of chilling stress under light (200 µmol photons m^-2^ s^-1^). Chilling-treated plants were dark-adapted for 30 min at 4 °C, while control plants were adapted at room temperature. The Pm values measured before chilling treatment were normalized to 1. Values are the mean ± SE, n = 3, biological replicates.

Next, the relationship between PSI photoinhibition by chilling stress in cucumber and PSII photoinhibition was investigated in more detail. Chilling stress was applied to cucumber plants under five light intensities, ranging from low light (80 µmol photons m^−2^ s^−1^) to high light (650 µmol photons m^−2^ s^−1^) to modulate the degree of PSII photoinhibition. As expected, higher light intensities induced stronger PSII photoinhibition, as indicated by a decrease in Fv/Fm below 0.2 under high light compared to around 0.5 under low light (Fig. 4A). In contrast, Pm dramatically decreased under all light intensities after chilling stress, around 20% under high light and around 10% under low light (Fig. 4A). Notably, PSI photoinhibition was more severe under lower light, suggesting a prolonged electron influx from PSII to PSI caused more severe PSI photoinhibition even in cucumber. When plotting Fv/Fm against Pm for each sample after chilling stress, a negative correlation was observed (correlation coefficient *r* = −0.68) (Fig. 4B), indicating that plants with greater PSII photoinhibition experienced less PSI photoinhibition.

**Figure 4.**
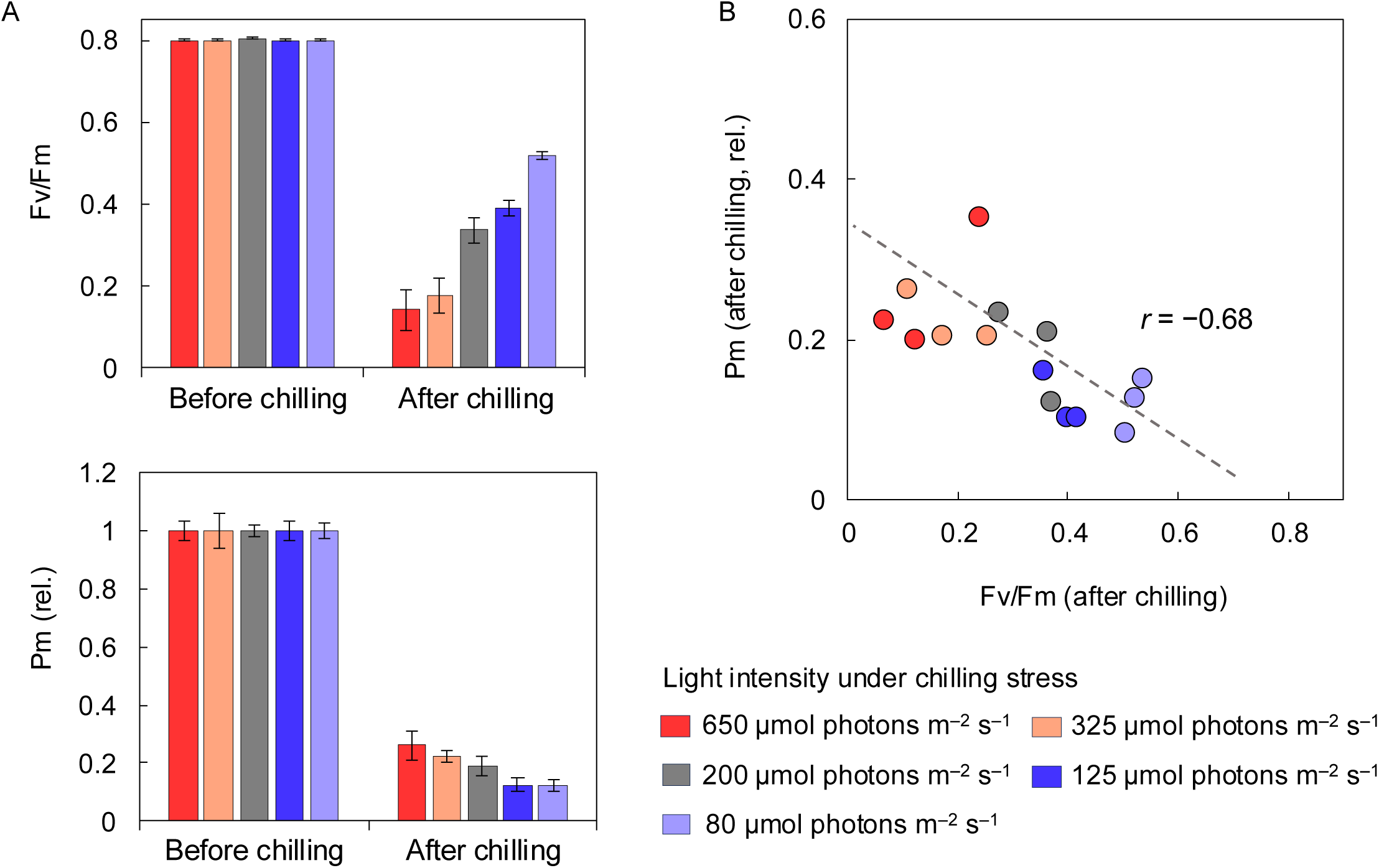
The relationship of PSII photoinhibition and PSI photoinhibition under chilling stress in cucumber. (A) Fv/Fm and Pm (rel.) before and after chilling stress for each light intensity. Cucumber leaf discs were floated in tap water and were exposed to chilling stress at 4 °C under light intensities ranging from 80 to 650 µmol photons m^-2^ s^-1^ for 20 h. At least 20 minutes of dark adaptation was performed before measuring Fv/Fm and Pm. The Pm values measured before chilling treatment were normalized to 1. (B) Correlation between Fv/Fm (measured after chilling treatment) and Pm (measured after chilling treatment). Each point represents biological replicates from 15 plant samples. The correlation coefficient *r* was −0.68. Values are the mean ± SE, n = 3, biological replicates.

### 3.3 Protective effect of PSII photoinhibition on PSI photoinhibition under fluctuating light

Next, we evaluated the protective effect of PSII photoinhibition on PSI photoinhibition under fluctuating light stress in *A. thaliana*. We applied fluctuating light stress to *A. thaliana* using repetitive short-pulse light (rSP), which selectively inhibits PSI (Sejima et al., 2014). In this experiment, non-chilled plants were used as controls, while plants subjected to long-chilling stress, which reduced Fv/Fm to approximately 0.4 (Supplementary Fig. S5), were used as samples suffering from PSII photoinhibition. As reported in previous studies, photo-oxidizable P700 decreased with increasing time of fluctuating light treatment in control plants (Fig. 5A–B) (Tsuyama and Kobayashi, 2009; Sejima et al., 2014; Zivcak et al., 2015; Takagi et al., 2017a; Shimakawa et al., 2019; Shimakawa and Miyake, 2019; Takagi et al., 2019; Ozaki et al., 2022; Takagi and Tani, 2023; Kono et al., 2025). In contrast, plants with severe PSII photoinhibition exhibited no decrease in active PSI even under fluctuating light; that is, PSI photoinhibition was markedly suppressed (Fig. 5A–B, and Supplementary Fig. S5). Y(II) decreased gradually in non-chilled plants but remained constant in plants after long-chilling stress (Fig. 5B). The residual activity of PSI after fluctuating light stress was reduced to 40% in non-chilled plants compared to before fluctuating light treatment, whereas PSI remained completely active in plants after long-chilling stress (Fig. 5C and Supplementary Fig. S5).

**Figure 5.**
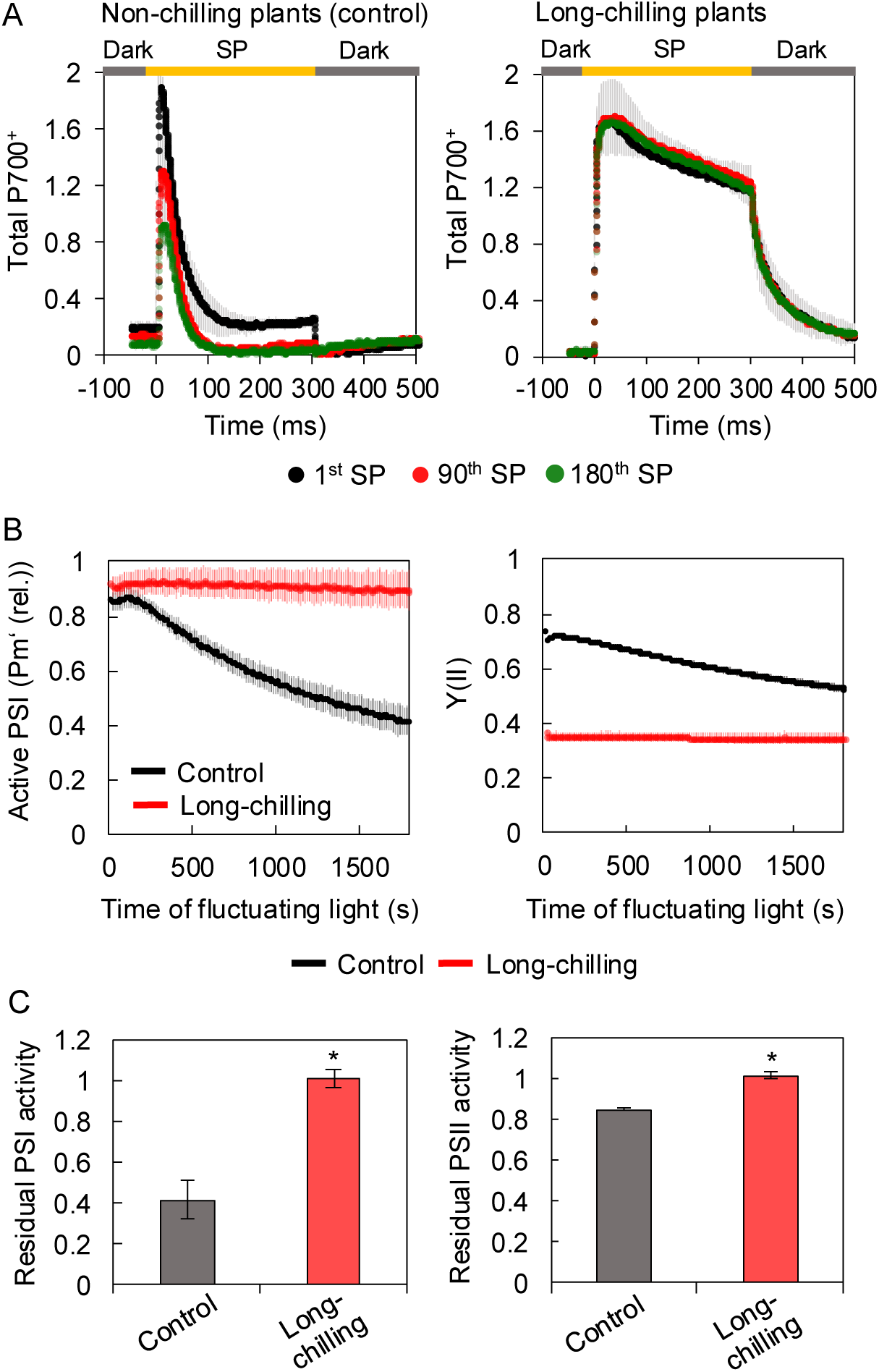
Effect of fluctuating light (rSP) treatment after long-chilling stress in *A. thaliana*. Fluctuating light treatment was applied to the long-chilled or non-chilled *A. thaliana* leaves using repetitive short-pulse (rSP) treatment. Leaves were exposed to short pulses (20000 µmol photons m^-2^ s^-1^, 300 ms) every 10 s for 30 min in the dark (a total of 180 SP). During the fluctuating light treatment, the leaf temperature was maintained at 23 °C. (A) Kinetics of P700^+^ during fluctuating light illumination. The data were obtained at the first SP (1^st^ SP), after 15 min (90^th^ SP), and after 30 min (180^th^ SP) during fluctuating light treatment. (B) Active PSI (Pm’(rel.)) and Y(II) during fluctuating light treatment. Active PSI was calculated as Pm’/Pm (relative Pm’) because P was not detected during the rSP treatment (Supplementary Fig. S5). Pm was measured before rSP treatment. (C) Residual PSI and PSII activity. The leaves were kept in the dark for 30 min after fluctuating light treatment, and residual activities of PSI and PSII were assessed by measuring Pm and Fv/Fm and normalizing the values of those measured before fluctuating light treatment to 1. Asterisks indicate statistically significant differences determined by Student’s t-test (*p* < 0.05). Values are the mean ± SE, n = 3, biological replicates.

The same experiment as Fig. 5 was carried out with plants of various Fv/Fm. *A. thaliana* with different degrees of PSII photoinhibition (Fv/Fm ranging from 0.2 to 0.8) were prepared by varying the duration of chilling stress. These plants were then subjected to fluctuating light treatment, during which the active PSI amount was calculated based on the decrease in Pm caused by fluctuating light treatment. Figure 6A shows the correlation between the degree of PSII photoinhibition before fluctuating light (Fv/Fm) and residual active PSI (Pm) after fluctuating light. Notably, the Pm after fluctuating light stress was strongly negatively correlated with Fv/Fm (correlation coefficient: *r* = −0.97) (Fig. 6A). That is, plants with lower Fv/Fm showed higher resistance to PSI photoinhibition by fluctuating light. The decline in active PSI during rSP treatment also exhibited a slope that correlated with the Fv/Fm values of the plants (Fig. 6B). These findings clearly demonstrate that PSI is protected from over-reduction by PSII photoinhibition, and susceptibility to PSI photoinhibition is strongly linked to Fv/Fm.

**Figure 6.**
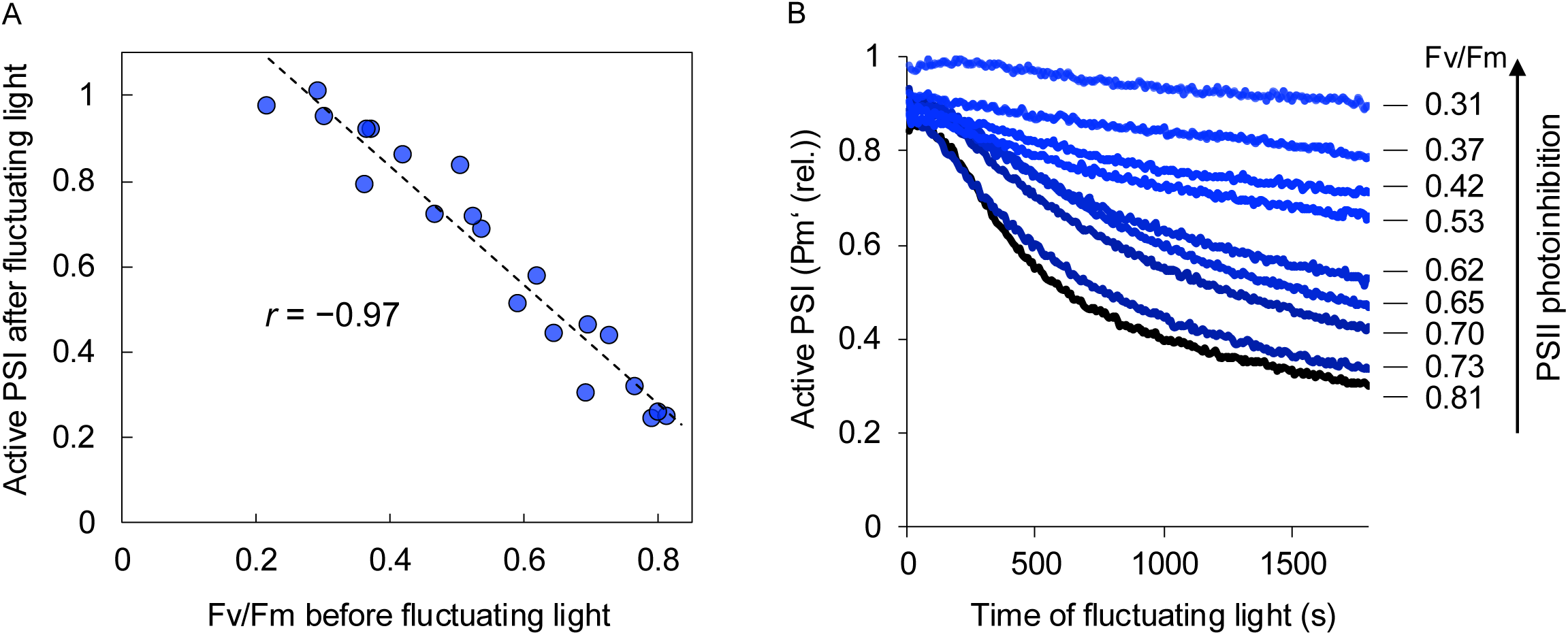
The relationship between PSII photoinhibition and PSI photoinhibition under fluctuating light in *A. thaliana*. *A. thaliana* WT with different degrees of PSII photoinhibition, indicated by Fv/Fm ranging from 0.2 to 0.8, were prepared by altering the treatment time of chilling stress. These plants were then subjected to fluctuating light (rSP) treatment at 23 °C, and the PSI photoinhibition was examined as the change in Pm. (A) Correlation between Fv/Fm (measured before fluctuating light treatment) and active PSI (residual Pm after rSP). Active PSI after fluctuating light was calculated by (Pm before rSP)/(Pm after rSP). Each point represents biological replicates from 21 plant samples. The correlation coefficient *r* was −0.97. (B) Decay kinetics of active PSI in representative plants during fluctuating light (rSP) treatment. Active PSI was calculated as Pm’/Pm (relative Pm’). PM was measured before rSP treatment. Fv/Fm values for each plant sample before rSP treatment are shown on the right edge of the graph.

### 3.4 Oxidation-reduction dynamics of Fe-S clusters in PSII photoinhibited plants

PSI photoinhibition results from ROS generated by the reaction of oxygen with reducing electron carriers downstream of P700 (Khorobrykh et al., 2020). We hypothesized that long-chilling-induced PSII photoinhibition in *A. thaliana* would mitigate the over-reduction of Fe-S clusters, thereby making PSI resistant to photoinhibition. To test this, *A. thaliana* plants subjected to long-chilling stress, whose P700 under SP is highly oxidized (Fig. 2 and Fig. 5A, right), were used to measure the oxidation-reduction levels of Fe-S clusters during SP following dark adaptation with Dual/KLAS-NIR. Unexpectedly, the Fe-S clusters were mostly reduced by SP in both control plants and plants exposed to long-chilling stress, and they remained predominantly in the reduced form throughout 300 ms of SP illumination (Fig. 7A). Given the depletion of electron flow from PSII after long-chilling stress (Fig. 1 and 2), Fe-S clusters must be reduced by charge separation of P700. Importantly, during the dark period after SP (after 300 ms), the reduced Fe-S clusters were re-oxidized more rapidly in plants subjected to long-chilling stress than in control plants (Fig. 7A). As mentioned above, the rate of re- oxidation of reduced Fe-S clusters after light illumination depends on the activation level of the electron efflux reactions from Fe-S clusters. Gas exchange measurements showed that the CBB cycle was largely suppressed after long-chilling stress, suggesting the proportion of electrons consumed by the downstream reactions would be low (Fig. 7B–C). Given that P700 was highly oxidized by PSII photoinhibition, this rapid re-oxidation of Fe-S clusters could be caused by an enhanced charge recombination reaction within PSI, which effectively prevents electron leakage to oxygen and PSI photoinhibition.

**Figure 7.**
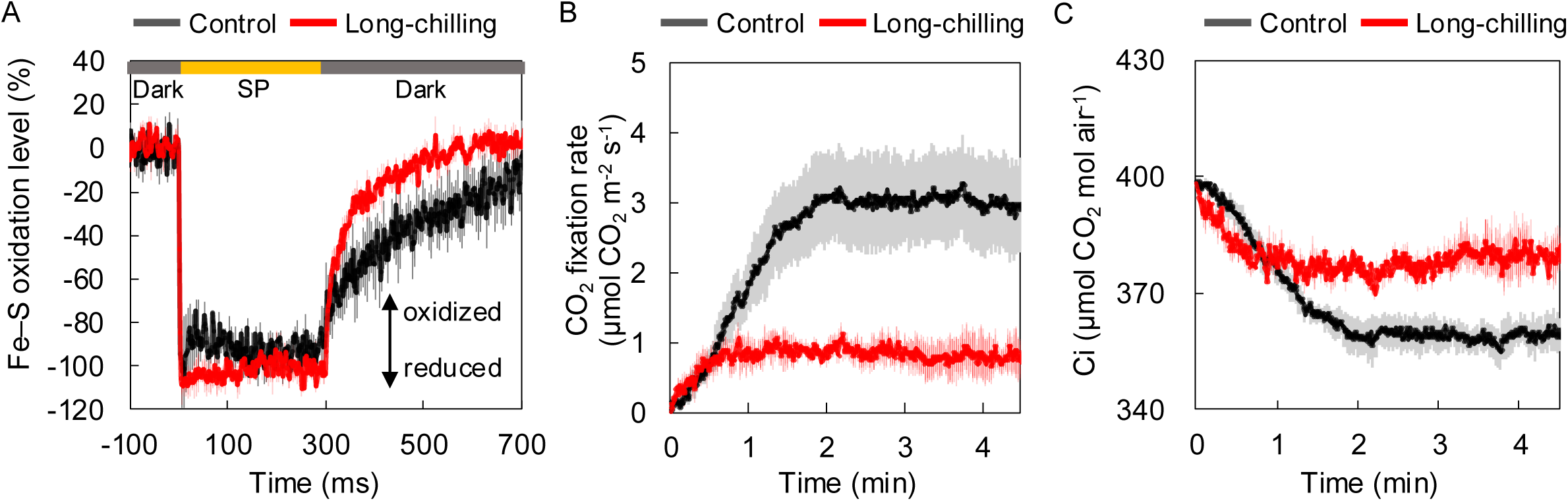
The redox changes of Fe-S clusters and CO_2_ fixation rate after long-chilling stress in *A. thaliana*. *A. thaliana* WT plants were grown for 4–6 weeks, and intact rosette leaves were analyzed before chilling stress (Control) and after long-chilling stress (Long-chilling). (A) Kinetics of reduced Fe-S clusters were measured at 23 °C under the illumination of SP (20000 µmol photons m^-2^ s^-1^ for 300 ms) after 20 min dark adaptation using Dual/KLAS-NIR. (B) CO_2_ fixation rate and (C) intercellular CO_2_ concentration (Ci) were determined by gas exchange analysis for 4–5 min after the onset of actinic light of 172 μmol photons m^−2^ s^−1^ after 20 min of dark adaptation at 25 °C. Values are the mean ± SE, n = 3, biological replicates.

## 4 Discussion

This study first clarified that PSII photoinhibition, induced by environmental stress, protects PSI in intact WT plants without using mutants or chemical treatments. Long-chilling stress caused severe PSII photoinhibition as indicated by the drastically decreased Fv/Fm and a significant decrease in electron influx into P700 in *A. thaliana* (Fig. 1 and 2). *A. thaliana* with decreased Fv/Fm exhibited higher tolerance against PSI photoinhibition (Fig. 5–6). Notably, susceptibility to PSI photoinhibition linearly correlated with Fv/Fm values (Fig. 6). In contrast, in cucumber, the suppression of PSI photoinhibition by PSII photoinhibition did not function effectively under chilling stress, leading to severe PSI damage (Fig. 3).

PSI photoinhibition is caused by ROS since it occurs only in the presence of oxygen (Sonoike and Terashima, 1994; Kono et al., 2014; Sejima et al., 2014; Takagi and Tani, 2023). In PSI, the electron transfer pathway follows P700→A_0_→A_1_→F_X_→F_A_/F_B_→Fd, where A_0_ is the primary acceptor chlorophyll *a*, A_1_ is the secondary acceptor phylloquinone, and F_X_, F_A_, and F_B_ are the iron-sulfur centers. All these components, including ^3^P700, have the potential to generate ROS (Kozuleva and Ivanov, 2010). In our experimental conditions, PSI photoinhibition—caused by an electron transfer from F_X_, F_A_, and F_B_ to O_2_ and the production of O_2_^•–^—could occur in plants during fluctuating light (rSP) treatment after long-chilling stress, given that Fe-S clusters were largely reduced during SP (Fig. 7) (Takahashi and Asada, 1988; Sonoike, 1996; Erling Tjus et al., 1998). However, PSI photoinhibition did not actually occur (Fig. 5). Consistently, O_2_^•–^, which could arise from reduced Fe-S clusters, was not increased by rSP treatment, at least in the soluble fraction as previously reported (Supplementary Fig. S6) (Mubarakshina et al., 2006). Our data support the previous ideas. First, the ROS formation during PSI photoinhibition by fluctuating light is mediated by carriers upstream of the Fe-S clusters, that is, O_2_^•–^ from phylloquinone A_1_ or ^1^O_2_ via the formation of ^3^P700 (Fig. 8) (Shuvalov et al., 1986; Rutherford et al., 2012; Cazzaniga et al., 2016; Kozuleva et al., 2021; Shimakawa et al., 2024); Second, PSI is resistant to photoinhibition as long as there is stable P700 charge separation (Tiwari et al., 2024). The prolonged lifetime of the reduced electron carriers in PSI increases their reactivity with oxygen, leading to ROS generation (Furutani et al., 2023). After long-chilling stress, the electrons in the Fe-S clusters were rapidly released (Fig. 7A), so that the electrons could not remain in the electron carriers long enough to react with oxygen. We propose that the charge recombination within PSI effectively scavenged electrons from reduced Fe-S clusters, since P700 was highly oxidized during SP (Fig. 5A). The stable back reaction from F_X_, F_A_, and F_B_ to phylloquinone molecules (A_1_) in A branches (A_1A_) leads to the formation of A_1A_^−^ rather than A_1_ in B branches (A_1B_^−^), thus avoiding the formation of ^3^P700 and allowing the excited P700 to return to the ground state (Fig.8) (Rutherford et al., 2012; Cherepanov et al., 2017; Degen and Johnson, 2024). Indeed, charge recombination followed by charge separation constitutes a local cycle in PSI under light illumination (Kou et al., 2015); it has been proposed to contribute to the difference between the total electron flux through PSI and the linear electron flux through both photosystems in leaves of low-light-grown plants exposed to high light stress. In addition to the proper charge recombination, plants have complex reactions downstream of Fd as electron escape pathways to prevent PSI photoinhibition. These reactions include ferredoxin- NADP^+^ reductase (FNR) to CBB cycle and photorespiration pathway, CEF, O_2_-dependent electron flow (water-water cycle, WWC), ferredoxin-thioredoxin reductase (FTR), and other metabolic pathways such as nitrogen assimilation, sulfur assimilation, chloroplast malate valve, and respiration (Fig. 8) (Schreiber, 2017; Nikkanen et al., 2018; Höhner et al., 2021; Krämer et al., 2024; Yamada et al., 2024). The effect of these diverse reactions on the role of electron scavenging from Fe-S needs to be further investigated.

**Figure 8.**
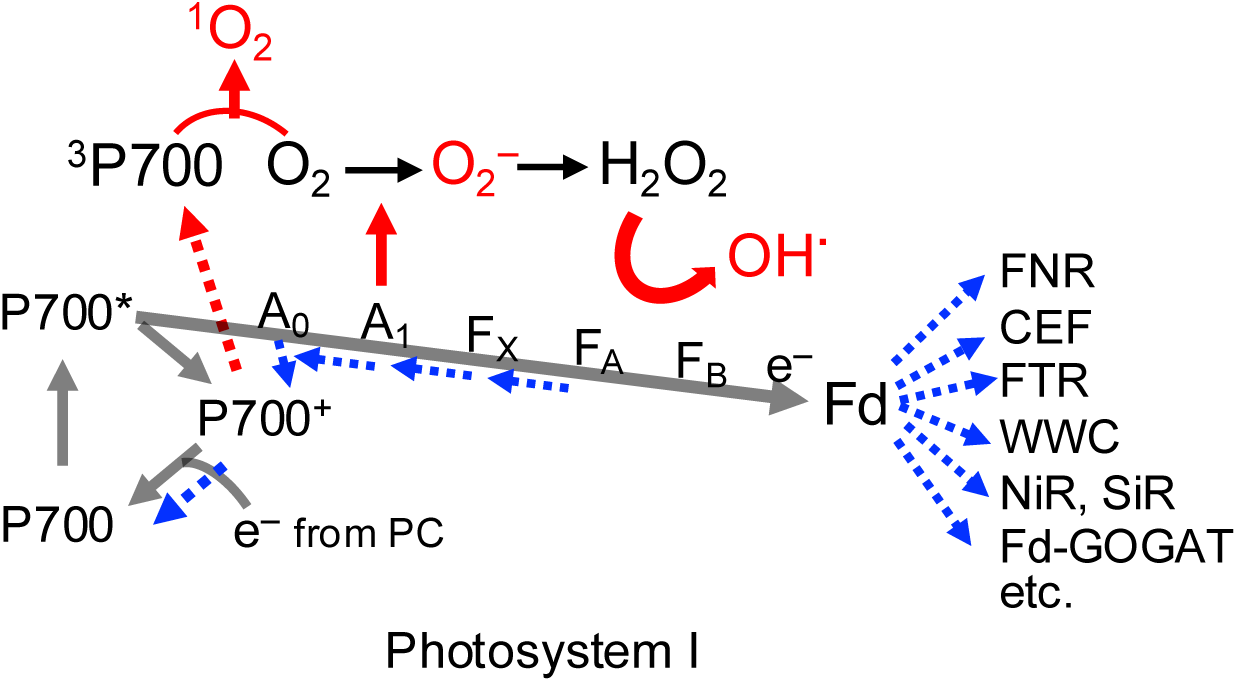
Scheme of electron flow and ROS generation in PSI under fluctuating light. In PSI, the electron transfer pathway follows P700→A_0_→A_1_→F_X_→F_A_/F_B_→Fd, where A_0_ is the primary acceptor chlorophyll *a*, A_1_ is the secondary acceptor phylloquinone, and F_X_, F_A_, and F_B_ are Fe-S clusters. In our view, if PSII is not properly down-regulated under environmental stress, ROS are generated either via direct electron transfer from phylloquinone to oxygen or through ^1^O_2_ formed from ^3^P700 (red arrow) under fluctuating light. Charge recombination from the B-branch during back reactions contributes to the formation of ^3^P700 (Rutherford et al., 2012). The generated ROS can subsequently react with Fe–S clusters to produce OH^•^. To suppress ROS production in PSI under fluctuating light, appropriate charge recombination reactions to P700^+^ and electron efflux downstream of Fd are considered important (blue arrow). Adequate P700^+^ accumulation by PSII photoinhibition enables the elimination of over-reduction downstream of PSI through charge recombination. Electron efflux downstream of Fd occurs via various pathways, including ferredoxin-NADP^+^ reductase (FNR) to the CBB cycle or photorespiratory pathway, CEF, ferredoxin-thioredoxin reductase (FTR), O_2_- dependent electron flow (WWC), and other metabolic pathways such as nitrogen assimilation, sulfur assimilation, chloroplast malate valve, respiration, etc. These diverse AEF pathways also collectively help in eliminating PSI over-reduction.

Under chilling stress, *A. thaliana* (chilling-tolerant plants) exhibited PSII photoinhibition-induced P700 oxidation, which suppressed PSI photoinhibition (Fig. 1–2). In contrast, cucumbers experienced PSI photoinhibition due to the insufficient downregulation of PSII activity (Fig. 3). In this case, a decrease in the re-oxidation rate of Fe-S clusters was observed (Supplementary Fig. S4), supporting the data of over-reduction of Fe-S and the inhibition of stable charge recombination in cucumber (Shimakawa et al., 2024; Takeuchi et al., in preparation). The cause of Fe-S over-reduction is known to be associated with the uncoupling of thylakoid membranes at low temperatures, leading to insufficient PSII downregulation by photosynthetic control (Peeler and Naylor, 1988; Terashima et al., 1991a, 1991b). Furthermore, our latest studies have shown that another factor in the over-reduction of Fe-S is the destabilization of the chloroplast NADH dehydrogenase-like complex (NDH) at low temperatures (Takeuchi et al., in preparation). On the other hand, more detailed studies are needed on the type of ROS produced by the over-reduction of Fe-S under chilling stress. One promising possibility is O_2_^•–^ generation from Fe-S clusters, which is supported by the idea that Fe-S clusters may be easier to access O_2_. Our data on fluctuating light did not support ROS generation at Fe-S clusters; however, it suggests that the inhibition sites of PSI and the sites of ROS production differ between chilling stress and fluctuating light stress (Shimakawa et al., 2024). Furthermore, wherever ROS occur, ROS in the thylakoid membrane are likely to produce OH^•^ via the Fenton reaction around unstable Fe- S clusters, leading to severe oxidative damage in PSI (Sonoike et al., 1995; Sonoike, 1996; Erling Tjus et al., 1998; Kılıç et al., 2023). On the other hand, the contribution of ^1^O_2_ to chilling-induced PSI photoinhibition has not been well discussed and should be discussed in the future. The detection of ^1^O_2_ around the PSI core requires ns-scale time resolution analysis, including ^3^P700 detection.

Under fluctuating light conditions, PSI photoinhibition can easily occur if electron acceptors other than the CBB cycle, such as FLV and CEF, are not functioning or if there are problems with ΔpH formation (Yamamoto et al., 2016; Kono et al., 2017; Wada et al., 2018; Wang et al., 2020; Luu Trinh et al., 2021; Rühle et al., 2021; Rodriguez-Heredia et al., 2022; Chaturvedi et al., 2024; Degen et al., 2024; Ermakova et al., 2024). Importantly, PSI was protected even under fluctuating light by decreasing PSII activity to approximately 40% through long-chilling stress (Fig. 5–6), again evidencing that PSII photoinhibition protects PSI in a manner unrelated to ΔpH.

Pulse amplitude modulation (PAM) analysis of photosynthetic activity involves SP illumination to monitor fluorescence yield and absorption changes of chlorophylls during SP, providing information about electron transport activity. In particular, understanding the information derived from the P700^+^ signal during SP in the dark is important, as changes in this signal provide insights into the P700 oxidation system and the PSI electron transfer processes. Previous studies have shown that the P700 redox cycle is driven even during 300 ms SP, varies with ΔpH, nutrient condition, or the presence of FLV, and provides various indications of electron influx into PSI and electron efflux from PSI (Ilík et al., 2017; Schreiber, 2017; Takagi et al., 2017b; Shimakawa and Miyake, 2018b; Shimakawa et al., 2019; Takagi et al., 2019; Nikkanen et al., 2023; Kuroki et al., 2024). In this study, it is newly proposed that the decay curve of P700^+^ during an SP after dark adaptation reflects electron influx from PSII to PSI (Fig. 2A); An increase in the decay rate of P700^+^ during SP reflects an increase in electron influx to PSI, whereas a decrease in the decay rate of P700^+^ reflects a decrease of electron influx. In this analysis, FR was applied before an SP to simultaneously measure the total amount of P700^+^ (Pm). The accumulation of P700^+^ during FR varied depending on the chilling treatment time (Fig. 2A). Various interpretations have been proposed for the P700^+^ accumulation during FR, such as reflecting AEF activity, including CEF (e.g., Joliot P. and Joliot A. 2005; Schreiber 2017), susceptibility to charge recombination (Kim et al. 2001), or reflecting damage in LHCI (Takagi et al., 2022). Although FR preferentially excites PSI, electron influx from PSII also occurs, making it difficult to discuss the interpretations based solely on FR in photoinhibited leaves (Chow and Hope 2004; Fan et al. 2016). One possible factor contributing to the differences in FR sensitivity during chilling stress is that, in the short-chilling treatment, AEF is likely activated (Supplementary Fig. S1), making the oxidation of P700 more likely to occur during FR. In contrast, in the long-chilling treatment, the decreased electron sink activity downstream of Fd (Fig. 7) or reduced energy transfer in LHCI are likely to reduce the sensitivity of FR.

So far, accurate determination of the effective quantum yield of PSI using SP under steady-state light conditions is challenging (Fan et al., 2016; Furutani et al., 2022). Y(I) merely shows the ratio of photo-oxidizable P700 by SP illumination to the total P700 content (Klughammer and Schreiber, 1994). Even within a few milliseconds of SP illumination, the P700 photo-reduction/oxidation cycle is driven to reach another state under the light intensity of SP illumination, depending on the oxidation level of both the donor and acceptor sides of P700 (Furutani et al., 2022). In this sense, Y(I) just reflects the balance between electron influx from the PSI donor side and efflux to the PSI acceptor side under SP. For example, the limitation of electron flow at the donor side of PSI leads to the rapid oxidation of P700 by SP illumination, which causes the overestimation of Y(I) (Furutani et al., 2022). It is also problematic to use Y(I) or ETR(I) under conditions where PSI photoinhibition can easily occur, such as under almost all environmental stresses (including chilling stress) or in mutants (e.g., *pgr5*, NDH lack of mutants). This is because, as in cucumber cold stress and other situations, if P700^+^ is significantly decreased under actinic light due to acceptor-side limitation, the apparent increase in P700^+^ during SP illumination occurs, leading to an overestimation of Y(I) (Takeuchi et al., 2022). Furthermore, under conditions where Pm changes significantly during measurement, current methods for assessing Y(I) consistently overestimate it due to the assumption that Pm remains constant. This problem has recently been addressed by (Grebe et al., 2024); PSI quantum yields assume a constant Pm and do not account for PSI photoinhibition, causing an overestimation of Y(I). To avoid overestimating Y(I) during PSI photoinhibition, they developed a new equation to calculate Y(I) that incorporates reductions in Pm due to PSI photoinhibition, demonstrating a truly balanced relationship between Y(II) and Y(I) under environmental stress. This means that the CEF activity, calculated by the gap between ETR(I) and ETR(II), is in fact hardly up-regulated in many studies under the condition of PSI photoinhibition (Grebe et al., 2024). We also emphasize that the current interpretations of Y(I) and ETR(I) are merely based on changes in the redox states of P700 under SP, and do not give any meaningful insight.

Long-chilling stress caused severe PSII photoinhibition in *A. thaliana* (Fig. 1). Adopting a simple model that considers the pool of PSII complexes as an ‘average’ population without splitting it into subpopulations differing in the oxidation state of Q_A_ or in the ability to perform stable charge separation (Kornyeyev and Hendrickson 2007), then Y(NO) determined as Fs/(Fm)PI, where (Fm)PI was measured after dark treatment following a chilling treatment that had induced photoinhibition (PI), is the sum of contributions from non-functional as well as functional PSII. That is, Y(NO) thus measured is an indicator of photoinhibition, as is Fv/Fm (Fig. 1). In contrast, care should be taken when interpreting qL in severely photoinhibited leaves. In this study, qL was calculated using a lake model as (Fm’– Fs)/(Fm’– Fó) × Fó/Fs. A low qL observed after long-chilling stress (Fig. 1) is typically interpreted as highly reduced Q_A_. However, Q_A_ would not actually be reduced since severely photoinhibited PSII complexes are incapable of Q_A_ photoreduction (Cleland et al. 1986). The observed low qL in Fig. 1 is primarily due to Fm′ being significantly reduced to a level comparable to Fs, as severe PSII photoinhibition suppresses PSII charge separation.

Overall, our results demonstrate that PSII photoinhibition induces P700 oxidation and protects PSI from photoinhibition. While a decline in Fv/Fm is often viewed as the primary negative consequence of low-temperature stress, it is important to recognize that plants decrease Fv/Fm as a protective mechanism to prevent PSI photoinhibition, which can offer positive effects as a survival strategy.

## Author contribution

Ko Takeuchi and Kentaro Ifuku designed the research. Ko Takeuchi and Shu Maekawa performed the experiments. Chikahiro Miyake provided the Dual/KLAS-NIR and LI-7000. Ko Takeuchi, Kentaro Ifuku, Chikahiro Miyake, Shintaro Harimoto, and Shu Maekawa analyzed data. Ko Takeuchi and Kentaro Ifuku wrote the manuscript.

## Data availability statement

Data sharing is not applicable to this article, as all newly created data is already contained within this article.

## Supporting information

Additional supporting information can be found online in the Supporting Information section at the end of this article.

## Supplementary data

**Supplementary Figure S1.**
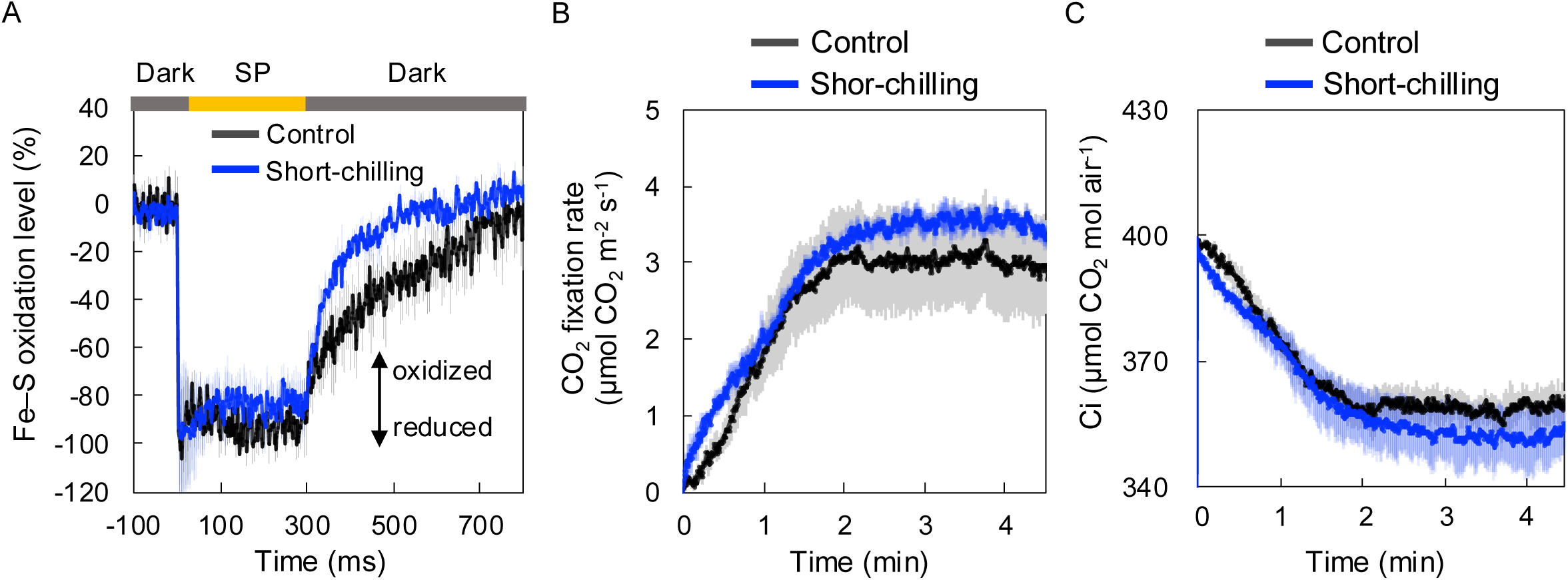
Responses downstream of P700 after short-chilling stress in *A. thaliana*. *A. thaliana* WT plants were grown for 4–6 weeks, and intact rosette leaves were analyzed before chilling stress (Control) and after 1 h of chilling stress (Short-chilling). (A) Kinetics of reduced Fe–S clusters were measured at 23 °C under the illumination of an SP (20000 μmol photons m-2 s-1 for 300 ms) after 20 min of dark adaptation using Dual/KLAS-NIR. (B) CO2 fixation rate and (C) intercellular CO2 concentration (Ci) were determined by gas exchange analysis for 4–5 min during the onset of actinic light of 172 μmol photons m−2 s−1 after 20 min of dark adaptation. Values are the mean ± SE, n = 3, biological replicates.

**Supplementary Figure S2.**
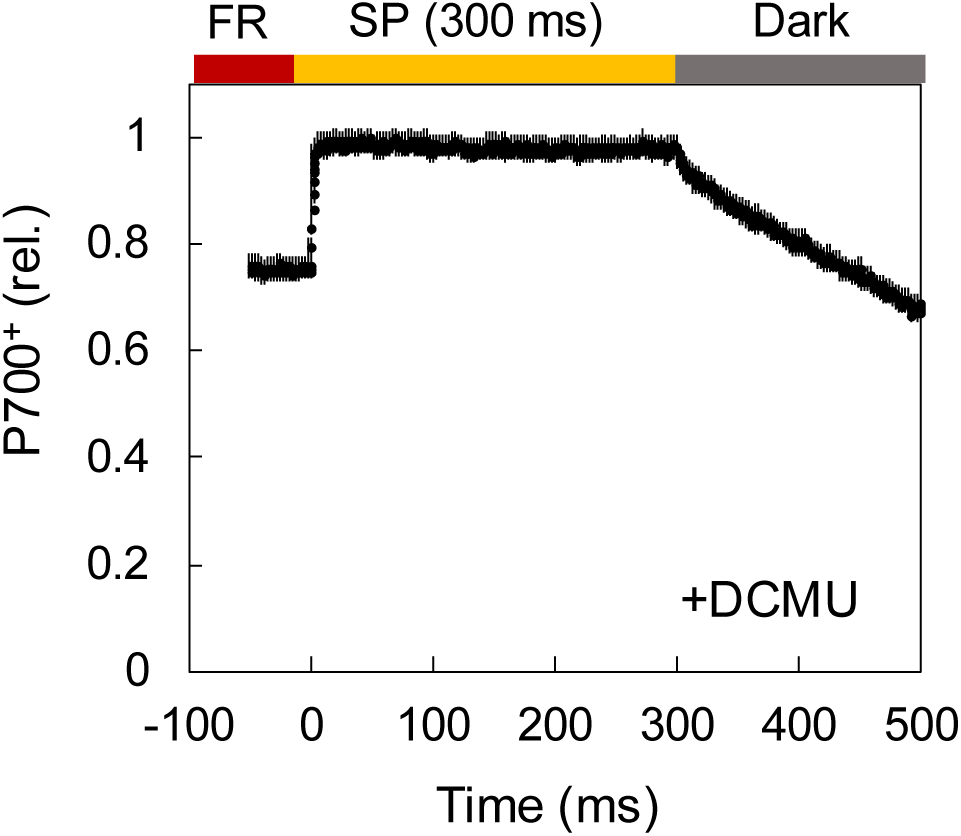
P700^+^ kinetics of DCMU-infiltrated leaves during SP. Leaves were vacuum-infiltrated with 50 µM DCMU for 1 min and dark-adapted for 60 min. Kinetics of P700^+^ during SP illumination were measured using the same method as for Pm determination. Leaves were first irradiated with far-red light for 10 s and then exposed to SP illumination of 20000 µmol photons m^-2^ s^-1^ for 300 ms under ambient conditions (23 °C). The obtained P700^+^ kinetics were normalized to a maximum value of 1.

**Supplementary Figure S3.**
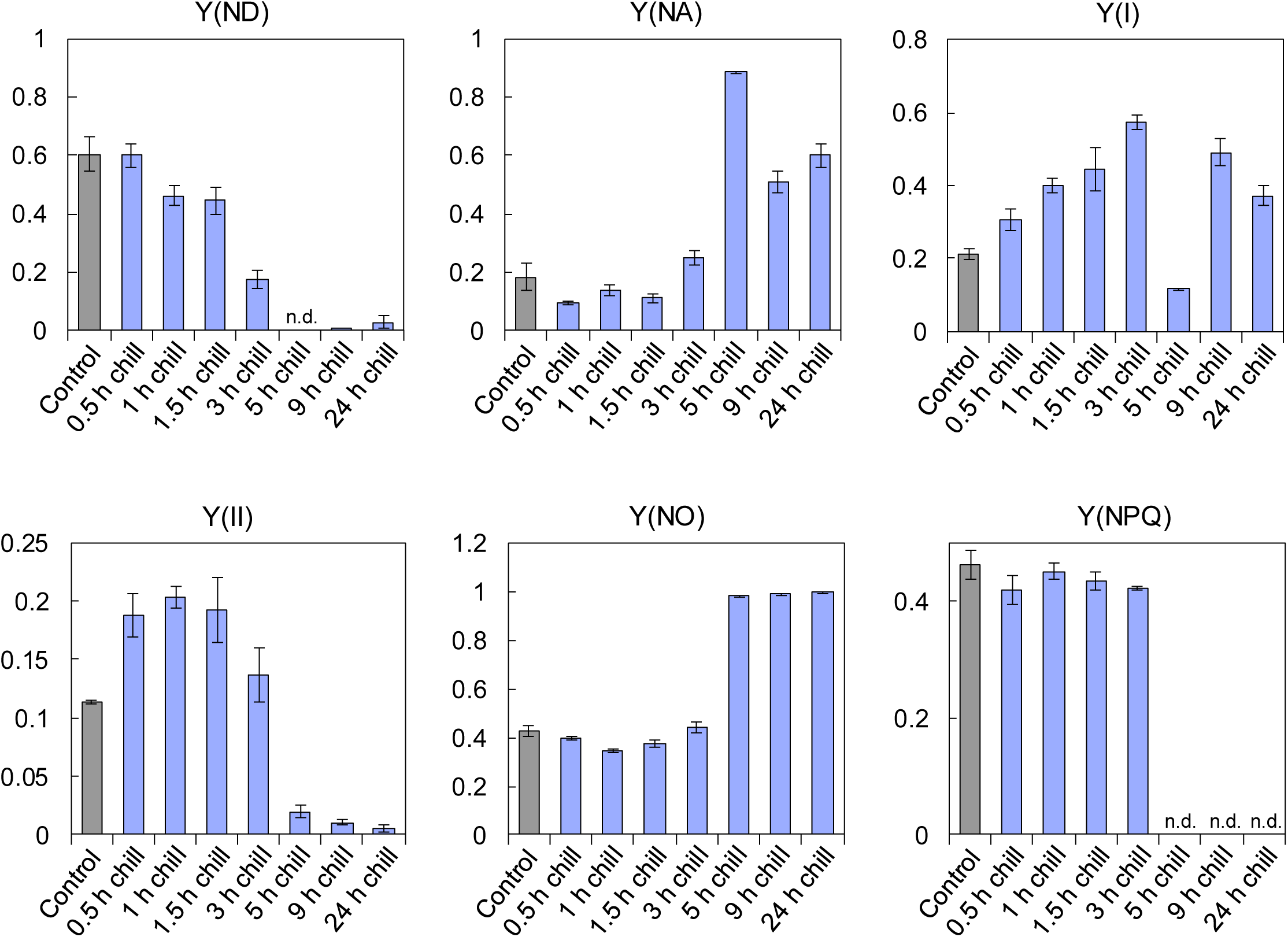
Effects of chilling stress on photosynthetic activity in cucumber. The 2–3-week-old cucumber plants were exposed to 0.5–24 h of chilling stress under middle light intensity (200 µmol photons m^-2^ s^-1^), and Chl fluorescence and P700 redox states were analyzed. Chilling-treated plants were dark-adapted for at least 20 min at 4 °C, while control plants were adapted at room temperature before measurements. Chl fluorescence and P700 redox states under the induction phase of photosynthesis were measured using Dual-PAM 100 at 23 °C after the illumination of actinic light (325 µmol photons m^−2^ s^−1^) for 5 min. Values are the mean ± SE, n = 3, biological replicates. ‘n.d.’ indicates no detectable value (0 value)

**Supplementary Figure S4.**
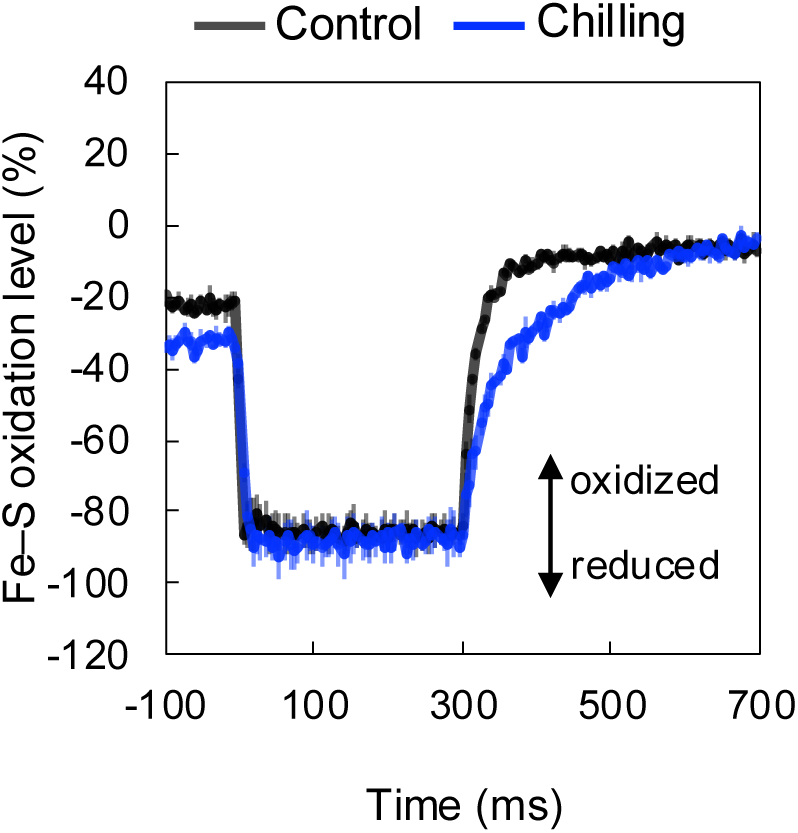
Fe-S redox kinetics before and after chilling stress in cucumber. Cucumber plants were grown for 2–3 weeks, and intact leaves were analyzed before chilling stress (Control), and after 5 h of chilling stress (Chilling). (A) Kinetics of reduced Fe–S clusters were measured under the illumination of an SP (20000 µmol photons m^-2^ s^-1^ for 300 ms) after 5 min of actinic light illumination at 23 °C using Dual/KLAS-NIR. During the dark period (300 ms to 700 ms) after SP illumination (0 ms to 300 ms), the reduced Fe-S clusters were re-oxidized. Values are the mean ± SE, n = 3, biological replicates.

**Supplementary Figure S5.**
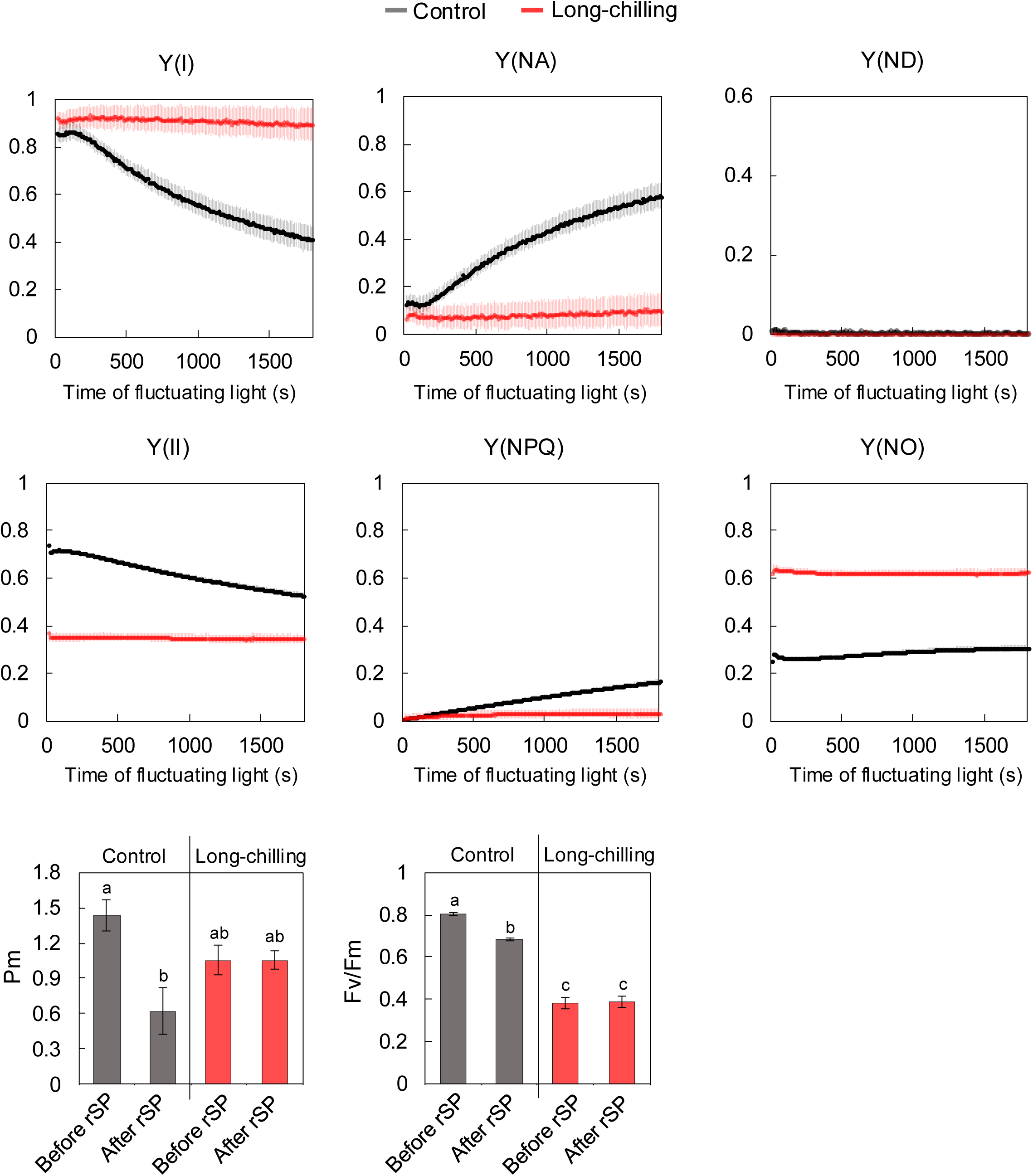
Photosynthetic parameters during fluctuating light (rSP) treatment before and after long- chilling stress. The plant samples and treatments were the same as those in Fig. 5. Fluctuating light treatment was applied to intact leaves (control and after long-chilling stress) using repetitive saturation pulse (rSP) treatment. Leaves were exposed to an SP (20000 µmol photons m^-2^ s^-1^, 300 ms) every 10 s for 30 min in the dark (a total of 180 SP). During the rSP treatment, the leaf temperature was maintained at 23 °C. Values are the mean ± SE, n = 3, biological replicates. Different letters indicate statistically significant differences by Tukey-Kramer’s multiple comparison tests after one-way ANOVA (*p* < 0.05).

**Supplementary Figure S6.**
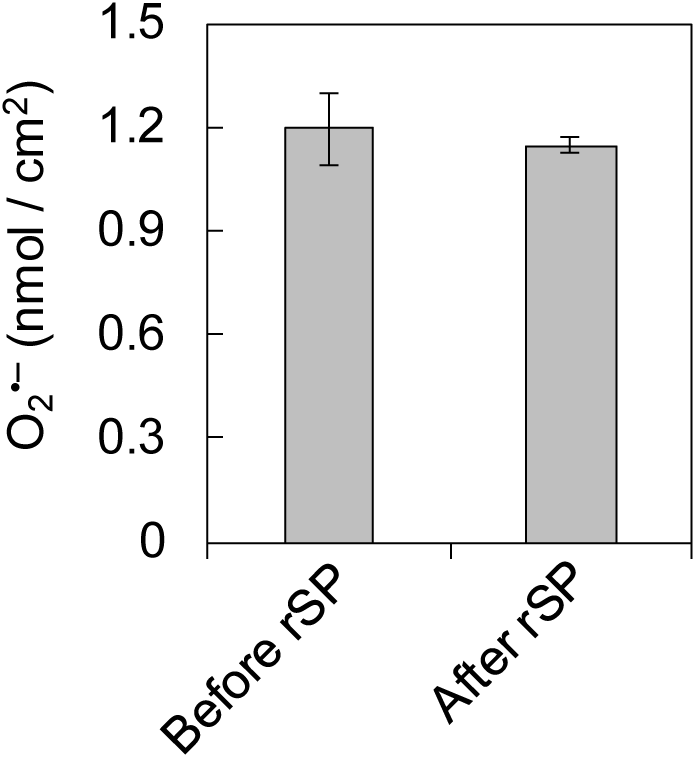
Accumulation of superoxide anion before and after rSP treatment. Fresh leaves (non-chilled) were harvested before and immediately after rSP treatment. The O_2_ ^•–^ content before and after rSP treatment was detected as described in materials and methods. Values are the mean ± SE, n = 3, biological replicates.

## Reference

Anderson JM (1992) Cytochrome *b*_6_*f* complex: dynamic molecular organization, function and acclimation. Photosynthesis Research 34: 341–357

Asada K (2006) Production and Scavenging of Reactive Oxygen Species in Chloroplasts and Their Functions. Plant Physiology 141: 391–396

Bag P, Shutova T, Shevela D, Lihavainen J, Nanda S, Ivanov AG, Messinger J, Jansson S (2023) Flavodiiron-mediated O_2_ photoreduction at photosystem I acceptor-side provides photoprotection to conifer thylakoids in early spring. Nature Communications 14: 3210

Baker NR, Harbinson J, Kramer DM (2007) Determining the limitations and regulation of photosynthetic energy transduction in leaves. Plant, Cell & Environment 30: 1107–1125

Barber J, Andersson B (1992) Too much of a good thing: light can be bad for photosynthesis. Trends in biochemical sciences 17: 61–66

Basso L, Sakoda K, Kobayashi R, Yamori W, Shikanai T (2022) Flavodiiron proteins enhance the rate of CO_2_ assimilation in Arabidopsis under fluctuating light intensity. Plant Physiology 189: 375– 387

Cazzaniga S, Bressan M, Carbonera D, Agostini A, Dall’Osto L (2016) Differential roles of carotenes and xanthophylls in photosystem I photoprotection. Biochemistry 55: 3636–3649

Chaturvedi AK, Dym O, Levin Y, Fluhr R (2024) PGR5-LIKE PHOTOSYNTHETIC PHENOTYPE1A redox states alleviate photoinhibition during changes in light intensity. Plant Physiology 194: 1059–1074

Chen S, Zheng Q, Qi Z, Ding J, Song X, Xia X (2024) Stress-induced delay of the IP rise of the fast chlorophyll *a* fluorescence transient in tomato. Scientia Horticulturae 326: 112741

Cherepanov DA, Milanovsky GE, Petrova AA, Tikhonov AN, Semenov AY (2017) Electron transfer through the acceptor side of photosystem I: Interaction with exogenous acceptors and molecular oxygen. Biochemistry Moscow 82: 1249–1268

Chow WS, Hope AB (2004) Electron fluxes through photosystem I in cucumber leaf discs probed by far-red light. Photosynthesis Research 81: 77–89

Cleland RE, Melis A, Neale PJ (1986) Mechanism of photoinhibition: photochemical reaction center inactivation in system II of chloroplasts. Photosynthesis Research 9: 79–88

Degen GE, Johnson MP (2024) Photosynthetic control at the cytochrome *b*_6_*f* complex. The Plant Cell 36: 4065–4079

Degen GE, Pastorelli F, Johnson MP (2024) Proton Gradient Regulation 5 is required to avoid photosynthetic oscillations during light transitions. Journal of Experimental Botany 75: 947–961

Dorion S, Ouellet JC, Rivoal J (2021) Glutathione metabolism in plants under stress: beyond reactive oxygen species detoxification. Metabolites 11: 641

Erling Tjus S, Lindberg Møller B, Vibe Scheller H (1998) Photosystem I is an early target of photoinhibition in barley illuminated at chilling temperatures. Plant Physiology 116: 755–764

Ermakova M, Woodford R, Fitzpatrick D, Nix SJ, Zwahlen SM, Farquhar GD, von Caemmerer S, Furbank RT (2024) Chloroplast NADH dehydrogenase-like complex-mediated cyclic electron flow is the main electron transport route in C_4_ bundle sheath cells. New Phytologist 243: 2187–2200

Fan D-Y, Fitzpatrick D, Oguchi R, Ma W, Kou J, Chow WS (2016) Obstacles in the quantification of the cyclic electron flux around Photosystem I in leaves of C3 plants. Photosynthesis Research 129: 239–251

Foyer C, Furbank R, Harbinson J, Horton P (1990) The mechanisms contributing to photosynthetic control of electron transport by carbon assimilation in leaves. Photosynthesis Research 25: 83–100

Foyer CH, Hanke G (2022) ROS production and signalling in chloroplasts: cornerstones and evolving concepts. The Plant Journal 111: 642–661

Furutani R, Makino A, Suzuki Y, Wada S, Shimakawa G, Miyake C (2020) Intrinsic fluctuations in transpiration induce photorespiration to oxidize P700 in photosystem I. Plants 9: 1761

Furutani R, Ohnishi M, Mori Y, Wada S, Miyake C (2022) The difficulty of estimating the electron transport rate at photosystem I. Journal of Plant Research 135: 565–577

Furutani R, Wada S, Ifuku K, Maekawa S, Miyake C (2023) Higher Reduced State of Fe/S-Signals, with the Suppressed Oxidation of P700, Causes PSI Inactivation in Arabidopsis thaliana. Antioxidants 12: 21

Gollan PJ, Grebe S, Roling L, Grimm B, Spetea C, Aro E (2023) Photosynthetic and transcriptome responses to fluctuating light in *Arabidopsis thylakoid* ion transport triple mutant. Plant Direct 7: e534

Grebe S, Porcar-Castell A, Riikonen A, Paakkarinen V, Aro E-M (2024) Accounting for photosystem I photoinhibition sheds new light on seasonal acclimation strategies of boreal conifers. Journal of Experimental Botany 75: 3973–3992

Hamada A, Tanaka Y, Ishikawa T, Maruta T (2023) Chloroplast dehydroascorbate reductase and glutathione cooperatively determine the capacity for ascorbate accumulation under photooxidative stress conditions. The Plant Journal 114: 68–82

Hanawa H, Ishizaki K, Nohira K, Takagi D, Shimakawa G, Sejima T, Shaku K, Makino A, Miyake C (2017) Land plants drive photorespiration as higher electron-sink: Comparative study of post-illumination transient O_2_-uptake rates from liverworts to angiosperms through ferns and gymnosperms. Physiologia Plantarum 161: 138–149

Hani U, Naranjo B, Shimakawa G, Espinasse C, Vanacker H, Sétif P, Rintamäki E, Issakidis-Bourguet E, Krieger-Liszkay A (2024) A complex and dynamic redox network regulates oxygen reduction at photosystem I in Arabidopsis. Plant Physiology kiae501

Havaux M, Davaud A (1994) Photoinhibition of photosynthesis in chilled potato leaves is not correlated with a loss of photosystem-II activity: preferential inactivation of photosystem I. Photosynthesis Research 40: 75–92

Heber U, Walker D (1992) Concerning a Dual Function of Coupled Cyclic Electron Transport in Leaves. Plant Physiology 100: 1621–1626

Helman Y, Barkan E, Eisenstadt D, Luz B, Kaplan A (2005) Fractionation of the three stable oxygen isotopes by oxygen-producing and oxygen-consuming reactions in photosynthetic organisms. Plant physiology 138: 2292–2298

Helman Y, Tchernov D, Reinhold L, Shibata M, Ogawa T, Schwarz R, Ohad I, Kaplan A (2003) Genes encoding A-type flavoproteins are essential for photoreduction of O_2_ in cyanobacteria. Current Biology 13: 230–235

Höhner R, Day PM, Zimmermann SE, Lopez LS, Krämer M, Giavalisco P, Correa Galvis V, Armbruster U, Schöttler MA, Jahns P (2021) Stromal NADH supplied by PHOSPHOGLYCERATE DEHYDROGENASE3 is crucial for photosynthetic performance. Plant Physiology 186: 142–167

Hope AB (2000) Electron transfers amongst cytochrome *f*, plastocyanin and photosystem I: kinetics and mechanisms. Biochimica et Biophysica Acta (BBA)-Bioenergetics 1456: 5–26

Huang W, Yang Y-J, Zhang S-B (2019) The role of water-water cycle in regulating the redox state of photosystem I under fluctuating light. Biochimica et Biophysica Acta (BBA)-Bioenergetics 1860: 383–390

Ifuku K, Endo T, Shikanai T, Aro E-M (2011) Structure of the chloroplast NADH dehydrogenase-like complex: nomenclature for nuclear-encoded subunits. Plant and Cell Physiology 52: 1560–1568

Ilík P, Pavlovič A, Kouřil R, Alboresi A, Morosinotto T, Allahverdiyeva Y, Aro E, Yamamoto H, Shikanai T (2017) Alternative electron transport mediated by flavodiiron proteins is operational in organisms from cyanobacteria up to gymnosperms. New Phytologist 214: 967–972

Inoue K, Fujii T, Yokoyama E, Matsuura K, Hiyama T, Sakurai H (1989) The photoinhibition site of photosystem I in isolated chloroplasts under extremely reducing conditions. Plant and cell physiology 30: 65–71

Ivanov AG, Allakhverdiev S, Hüner N, Murata N (2012a) Genetic decrease in fatty acid unsaturation of phosphatidylglycerol increased photoinhibition of photosystem I at low temperature in tobacco leaves. Biochimica et Biophysica Acta (BBA)-Bioenergetics 1817: 1374–1379

Ivanov AG, Morgan RM, Gray GR, Velitchkova MY, Hüner N (1998) Temperature/light dependent development of selective resistance to photoinhibition of photosystem I. FEBS letters 430: 288–292

Ivanov AG, Rosso D, Savitch L, Stachula P, Rosembert M, Oquist G, Hurry V, Hüner N (2012b) Implications of alternative electron sinks in increased resistance of PSII and PSI photochemistry to high light stress in cold-acclimated *Arabidopsis thaliana*. Photosynthesis research 113: 191–206

Joliot P, Joliot A (2005) Quantification of cyclic and linear flows in plants. Proceedings of the National Academy of Sciences 102: 4913–4918

Kadota K, Furutani R, Makino A, Suzuki Y, Wada S, Miyake C (2019) Oxidation of P700 induces alternative electron flow in photosystem I in wheat leaves. Plants 8: 152

Kale R, Hebert AE, Frankel LK, Sallans L, Bricker TM, Pospíšil P (2017) Amino acid oxidation of the D1 and D2 proteins by oxygen radicals during photoinhibition of Photosystem II. Proceedings of the National Academy of Sciences, USA 114: 2988–2993

Kanazawa A, Ostendorf E, Kohzuma K, Hoh D, Strand DD, Sato-Cruz M, Savage L, Cruz JA, Fisher N, Froehlich JE (2017) Chloroplast ATP synthase modulation of the thylakoid proton motive force: implications for photosystem I and photosystem II photoprotection. Frontiers in Plant Science 8: 719

Khorobrykh S, Havurinne V, Mattila H, Tyystjärvi E (2020) Oxygen and ROS in photosynthesis. Plants 9: 91

Kim J, Kim S, Cho SH, Chow WS, Lee C (2005) Photosystem I acceptor side limitation is a prerequisite for the reversible decrease in the maximum extent of P700 oxidation after short-term chilling in the light in four plant species with different chilling sensitivities. Physiologia Plantarum 123: 100–107

Kim S, Lee C, Hope A, Chow WS (2001) Inhibition of photosystems I and II and enhanced back flow of photosystem I electrons in cucumber leaf discs chilled in the light. Plant and Cell Physiology 42: 842–848

Kılıç M, Käpylä V, Gollan PJ, Aro E-M, Rintamäki E (2023) PSI Photoinhibition and Changing CO_2_ Levels Initiate Retrograde Signals to Modify Nuclear Gene Expression. Antioxidants 12: 1902

Klughammer C, Schreiber U (2008) Saturation Pulse method for assessment of energy conversion in PS I. PAM Application Notes 1: 11–14

Klughammer C, Schreiber U (1994) An improved method, using saturating light pulses, for the determination of photosystem I quantum yield via P700^+^-absorbance changes at 830 nm. Planta 192: 261–268

Klughammer C, Schreiber U (2016) Deconvolution of ferredoxin, plastocyanin, and P700 transmittance changes in intact leaves with a new type of kinetic LED array spectrophotometer. Photosynthesis Research 128: 195–214

Kohzuma K, Cruz JA, Akashi K, Hoshiyasu S, Munekage YN, Yokota A, Kramer DM (2009) The long-term responses of the photosynthetic proton circuit to drought. Plant, cell & environment 32: 209–219

Kono M, Kodama H, Tanigawa K, Terashima I, Yamori W (2025) Cyt *b*_6_/*f* complex fine-tunes PSI stability and photosynthetic capacity under fluctuating light. BioRxiv, 10.1101/2025.02.16.638563

Kono M, Noguchi K, Terashima I (2014) Roles of the cyclic electron flow around PSI (CEF-PSI) and O_2_-dependent alternative pathways in regulation of the photosynthetic electron flow in short-term fluctuating light in *Arabidopsis thaliana*. Plant and Cell Physiology 55: 990–1004

Kono M, Oguchi R, Terashima I (2022) Photoinhibition of PSI and PSII in Nature and in the Laboratory: Ecological Approaches. Progress in Botany 84: 241–292

Kono M, Yamori W, Suzuki Y, Terashima I (2017) Photoprotection of PSI by far-red light against the fluctuating light-induced photoinhibition in *Arabidopsis thaliana* and field-grown plants. Plant and Cell Physiology 58: 35–45

Kornyeyev D, Hendrickson L (2007) Research note: Energy partitioning in photosystem II complexes subjected to photoinhibitory treatment. Functional Plant Biology 34: 214–220

Kou J, Takahashi S, Fan D-Y, Badger MR, Chow WS (2015) Partially dissecting the steady-state electron fluxes in Photosystem I in wild-type and *pgr5* and *ndh* mutants of *Arabidopsis*. Frontiers in Plant Science 6: 758

Kozuleva M, Petrova A, Milrad Y, Semenov A, Ivanov B, Redding KE, Yacoby I (2021) Phylloquinone is the principal Mehler reaction site within photosystem I in high light. Plant Physiology 186: 1848–1858

Kozuleva MA, Ivanov BN (2010) Evaluation of the participation of ferredoxin in oxygen reduction in the photosynthetic electron transport chain of isolated pea thylakoids. Photosynthesis Research 105: 51–61

Kozuleva MA, Petrova AA, Mamedov MD, Semenov AY, Ivanov BN (2014) O_2_ reduction by photosystem I involves phylloquinone under steady-state illumination. FEBS letters 588: 4364–4368

Krämer M, Blanco NE, Penzler J-F, Davis GA, Brandt B, Leister D, Kunz H-H (2024) Cyclic electron flow compensates loss of PGDH3 and concomitant stromal NADH reduction. Scientific Reports 14: 29274

Krieger-Liszkay A (2005) Singlet oxygen production in photosynthesis. Journal of Experimental Botany 56: 337–346

Kumar A, Prasad A, Sedlářová M, Kale R, Frankel LK, Sallans L, Bricker TM, Pospíšil P (2021) Tocopherol controls D1 amino acid oxidation by oxygen radicals in Photosystem II. Proceedings of the National Academy of Sciences, USA. 118: e2019246118

Kuroki S, Ohnishi M, Furutani R, Tsuru K, Miyake C (2024) Nondestructive diagnosis of mineral deficiencies in wheat leaves by one-or two-shot saturation pulse measurement of chlorophyll fluorescence and P700^+^ absorbance with machine learning. Smart Agricultural Technology 9: 10058

Li J, Zhang H, Yue D, Chen S, Yin Y, Zheng C, Chen Y (2024) Endogenous serotonin induced by cold acclimation increases cold tolerance by reshaping the MEL/ROS/RNS redox network in *Kandelia obovata*. Journal of Forestry Research 35: 1–13

Li L, Aro E-M, Millar AH (2018) Mechanisms of photodamage and protein turnover in photoinhibition. Trends in Plant Science 23: 667–676

Lima-Melo Y, Gollan PJ, Tikkanen M, Silveira JA, Aro E (2019) Consequences of photosystem-I damage and repair on photosynthesis and carbon use in *Arabidopsis thaliana*. The Plant Journal 97: 1061–1072

Luu Trinh MD, Miyazaki D, Ono S, Nomata J, Kono M, Mino H, Niwa T, Okegawa Y, Motohashi K, Taguchi H, et al (2021) The evolutionary conserved iron-sulfur protein TCR controls P700 oxidation in photosystem I. iScience 24: 102059

Maekawa S, Ohnishi M, Wada S, Ifuku K, Miyake C (2024) Enhanced Reduction of Ferredoxin in PGR5-Deficient Mutant of *Arabidopsis thaliana* Stimulated Ferredoxin-Dependent Cyclic Electron Flow around Photosystem I. International Journal of Molecular Sciences 25: 2677

Mehler AH (1951) Studies on reactions of illuminated chloroplasts: I. Mechanism of the reduction of oxygen and other hill reagents. Archives of Biochemistry and Biophysics 33: 65–77

Melis A (1999) Photosystem-II damage and repair cycle in chloroplasts: what modulates the rate of photodamage *in vivo*? Trends in Plant Science 4: 130–135

Messant M, Hani U, Lai T, Wilson A, Shimakawa G, Krieger-Liszkay A (2024) Plastid terminal oxidase (PTOX) protects photosystem I and not photosystem II against photoinhibition in *Arabidopsis thaliana* and *Marchantia polymorpha*. The Plant Journal 117: 669–678

Miyake C (2020) Molecular mechanism of oxidation of P700 and suppression of ROS production in photosystem I in response to electron-sink limitations in C3 plants. Antioxidants 9: 230

Miyake C (2010) Alternative electron flows (water–water cycle and cyclic electron flow around PSI) in photosynthesis: molecular mechanisms and physiological functions. Plant and Cell Physiology 51: 1951–1963

Miyake C, Amako K, Shiraishi N, Sugimoto T (2009) Acclimation of tobacco leaves to high light intensity drives the plastoquinone oxidation system—relationship among the fraction of open PSII centers, non-photochemical quenching of Chl fluorescence and the maximum quantum yield of PSII in the dark. Plant and Cell Physiology 50: 730–743

Miyata K, Noguchi K, Terashima I (2012) Cost and benefit of the repair of photodamaged photosystem II in spinach leaves: roles of acclimation to growth light. Photosynthesis Research 113: 165–180

Mubarakshina M, Khorobrykh S, Ivanov B (2006) Oxygen reduction in chloroplast thylakoids results in production of hydrogen peroxide inside the membrane. Biochimica et Biophysica Acta (BBA)-Bioenergetics 1757: 1496–1503

Murata N, Takahashi S, Nishiyama Y, Allakhverdiev SI (2007) Photoinhibition of photosystem II under environmental stress. Biochimica et Biophysica Acta (BBA)-Bioenergetics 1767: 414–421

Nakano R, Ishida H, Kobayashi M, Makino A, Mae T (2010) Biochemical changes associated with *in vivo* RbcL fragmentation by reactive oxygen species under chilling-light conditions. Plant Biology 12: 35–45

Nakano R, Ishida H, Makino A, Mae T (2006) In vivo fragmentation of the large subunit of ribulose- 1, 5-bisphosphate carboxylase by reactive oxygen species in an intact leaf of cucumber under chilling-light conditions. Plant and Cell Physiology 47: 270–276

Napaumpaiporn P, Ogawa T, Sonoike K, Nishiyama Y (2024) Improved capacity for the repair of photosystem II via reinforcement of the translational and antioxidation systems in *Synechocystis* sp. PCC 6803. The Plant Journal 117: 1165–1178

Nikkanen L, Toivola J, Trotta A, Diaz MG, Tikkanen M, Aro E, Rintamäki E (2018) Regulation of cyclic electron flow by chloroplast NADPH-dependent thioredoxin system. Plant Direct 2: 1–24

Nikkanen L, Vakal S, Santana-Sánchez A, Hubáček M, Konert G, Wang Y, Boehm M, Gutekunst K, Salminen TA, Allahverdiyeva Y (2023) Flavodiiron proteins associate pH-dependently with the thylakoid membrane for ferredoxin-1 powered O_2_ photoreduction. BioRxiv 10.1101/2023.05.19.541409

Nishio JN, Whitmarsh J (1993) Dissipation of the proton electrochemical potential in intact chloroplasts (II. The pH gradient monitored by cytochrome f reduction kinetics). Plant Physiology 101: 89–96

Nishiyama Y, Allakhverdiev SI, Murata N (2006) A new paradigm for the action of reactive oxygen species in the photoinhibition of photosystem II. Biochimica et Biophysica Acta (BBA)-Bioenergetics 1757: 742–749

Nishiyama Y, Yamamoto H, Allakhverdiev SI, Inaba M, Yokota A, Murata N (2001) Oxidative stress inhibits the repair of photodamage to the photosynthetic machinery. The EMBO Journal 20: 5587–5594

Obara A, Ogawa M, Oyama Y, Suzuki Y, Kono M (2022) Effects of high irradiance and low water temperature on photoinhibition and repair of photosystems in Marimo (*Aegagropila linnaei*) in Lake Akan, Japan. International Journal of Molecular Sciences 24: 60

Ohnishi M, Maekawa S, Wada S, Ifuku K, Miyake C (2023) Evaluating the Oxidation Rate of Reduced Ferredoxin in *Arabidopsis thaliana* independent of photosynthetic linear electron flow: Plausible activity of ferredoxin-dependent cyclic electron flow around photosystem I. International Journal of Molecular Sciences 24: 12145

Oxborough K, Baker NR (1997) Resolving chlorophyll *a* fluorescence images of photosynthetic efficiency into photochemical and non-photochemical components – calculation of *qP* and *Fv*’/*Fm*’ without measuring *Fo*. Photosynthesis research 54: 135–142

Ozaki H, Takagi D, Mizokami Y, Tokida T, Nakamura H, Sakai H, Hasegawa T, Noguchi K (2022) Low N level increases the susceptibility of PSI to photoinhibition induced by short repetitive flashes in leaves of different rice varieties. Physiologia Plantarum 174: e13644

Ozawa S-I, Zhang G, Sakamoto W (2024) Dysfunction of Chloroplast Protease Activity Mitigates *pgr5* Phenotype in the Green Algae *Chlamydomonas reinhardtii*. Plants 13: 606

Peeler TC, Naylor AW (1988) A comparison of the effects of chilling on thylakoid electron transfer in pea (*Pisum sativum* L.) and cucumber (*Cucumis sativus* L.). Plant Physiology 86: 147–151

Petronek MS, Spitz DR, Allen BG (2021) Iron–sulfur cluster biogenesis as a critical target in cancer. Antioxidants 10: 1458

Pospíšil P, Yamamoto Y (2017) Damage to photosystem II by lipid peroxidation products. Biochimica et Biophysica Acta (BBA)-General Subjects 1861: 457–466

Powles SB (1984) Photoinhibition of photosynthesis induced by visible light. Annual Review of Plant Physiology 35: 15–44

Rodriguez-Heredia M, Saccon F, Wilson S, Finazzi G, Ruban AV, Hanke GT (2022) Protection of photosystem I during sudden light stress depends on ferredoxin: NADP (H) reductase abundance and interactions. Plant Physiology 188: 1028–1042

Rühle T, Dann M, Reiter B, Schünemann D, Naranjo B, Penzler J-F, Kleine T, Leister D (2021) PGRL2 triggers degradation of PGR5 in the absence of PGRL1. Nature Communications 12: 3941

Rutherford AW, Osyczka A, Rappaport F (2012) Back-reactions, short-circuits, leaks and other energy wasteful reactions in biological electron transfer: redox tuning to survive life in O_2_. FEBS letters 586: 603–616

Sacksteder CA, Kramer DM (2000) Dark-interval relaxation kinetics (DIRK) of absorbance changes as a quantitative probe of steady-state electron transfer. Photosynthesis Research 66: 145–158

Sadiq M, Akram NA, Ashraf M, Al-Qurainy F, Ahmad P (2019) Alpha-tocopherol-induced regulation of growth and metabolism in plants under non-stress and stress conditions. Journal of Plant Growth Regulation 38: 1325–1340

Schnettger B, Critchley C, Santore U, Graf M, Krause G (1994) Relationship between photoinhibition of photosynthesis, D1 protein turnover and chloroplast structure: effects of protein synthesis inhibitors. Plant, Cell & Environment 17: 55–64

Schöttler MA, Tóth SZ (2014) Photosynthetic complex stoichiometry dynamics in higher plants: environmental acclimation and photosynthetic flux control. Frontiers in Plant Science 5: 188

Schöttler MA, Tóth SZ, Boulouis A, Kahlau S (2015) Photosynthetic complex stoichiometry dynamics in higher plants: biogenesis, function, and turnover of ATP synthase and the cytochrome *b*_6_*f* complex. Journal of Experimental Botany 66: 2373–2400

Schreiber U (2017) Redox changes of ferredoxin, P700, and plastocyanin measured simultaneously in intact leaves. Photosynthesis Research 134: 343–360

Sejima T, Takagi D, Fukayama H, Makino A, Miyake C (2014) Repetitive short-pulse light mainly inactivates photosystem I in sunflower leaves. Plant and Cell Physiology 55: 1184–1193

Shaku K, Shimakawa G, Hashiguchi M, Miyake C (2016) Reduction-induced suppression of electron flow (RISE) in the photosynthetic electron transport system of *Synechococcus elongatus PCC* 7942. Plant and Cell Physiology 57: 1443–1453

Shikanai T (2007) Cyclic electron transport around photosystem I: genetic approaches. Annual Review of Plant Biology 58: 199–217

Shikanai T (2023) Molecular genetic dissection of the regulatory network of proton motive force in chloroplasts. Plant and Cell Physiology 65: 537–550

Shimakawa G, Iwamoto T, Mabuchi T, Saito R, Yamamoto H, Amako K, Sugimoto T, Makino A, Miyake C (2013) Acrolein, an α, β-unsaturated carbonyl, inhibits both growth and PSII activity in the cyanobacterium *Synechocystis* sp. PCC 6803. Bioscience, Biotechnology, and Biochemistry 77: 1655–1660

Shimakawa G, Miyake C (2018a) Oxidation of P700 Ensures Robust Photosynthesis. Frontiers in Plant Science 9: 1617

Shimakawa G, Miyake C (2019) What quantity of photosystem I is optimum for safe photosynthesis? Plant Physiology 179: 1479–1485

Shimakawa G, Miyake C (2018b) Respiratory terminal oxidases alleviate photo-oxidative damage in photosystem I during repetitive short-pulse illumination in *Synechocystis* sp. PCC 6803. Photosynthesis Research 137: 241–250

Shimakawa G, Müller P, Miyake C, Krieger-Liszkay A, Sétif P (2024) Photo-oxidative damage of photosystem I by repetitive flashes and chilling stress in cucumber leaves. Biochimica et Biophysica Acta (BBA)-Bioenergetics 149490

Shimakawa G, Murakami A, Niwa K, Matsuda Y, Wada A, Miyake C (2019) Comparative analysis of strategies to prepare electron sinks in aquatic photoautotrophs. Photosynthesis Research 139: 401–411

Shuvalov V, Nuijs A, Van Gorkom H, Smit H, Duysens L (1986) Picosecond absorbance changes upon selective excitation of the primary electron donor P-700 in photosystem I. Biochimica et Biophysica Acta (BBA)-Bioenergetics 850: 319–323

Sonoike K (2011) Photoinhibition of photosystem I. Physiologia Plantarum 142: 56–64

Sonoike K (1996) Photoinhibition of photosystem I: Its physiological significance in the chilling sensitivity of plants. Plant and Cell Physiology 37: 239–247

Sonoike K, Terashima I (1994) Mechanism of photosystem-I photoinhibition in leaves of *Cucumis sativus* L. Planta 194: 287–293

Sonoike K, Terashima I, Iwaki M, Itoh S (1995) Destruction of photosystem I iron-sulfur centers in leaves of *Cucumis sativus* L. by weak illumination at chilling temperatures. FEBS letters 362: 235– 238

Storti M, Puggioni MP, Segalla A, Morosinotto T, Alboresi A (2020) The chloroplast NADH dehydrogenase-like complex influences the photosynthetic activity of the moss *Physcomitrella patens*. Journal of Experimental Botany 71: 5538–5548

Sun H, Shi Q, Liu N-Y, Zhang S-B, Huang W (2023) Drought stress delays photosynthetic induction and accelerates photoinhibition under short-term fluctuating light in tomato. Plant Physiology and Biochemistry 196: 152–161

Suorsa M, Järvi S, Grieco M, Nurmi M, Pietrzykowska M, Rantala M, Kangasjärvi S, Paakkarinen V, Tikkanen M, Jansson S (2012) PROTON GRADIENT REGULATION5 is essential for proper acclimation of *Arabidopsis* photosystem I to naturally and artificially fluctuating light conditions. The Plant Cell 24: 2934–2948

Suzuki Y, Nagao K, Takahashi Y, Miyake C, Makino A (2021) Oxidation of the reaction center chlorophyll of photosystem I is induced via close cooperation of photosystems II and I with progress of drought stress in soybean seedlings. Soil Science and Plant Nutrition 67: 662–669

Tachibana R, Abe S, Marugami M, Yamagami A, Akema R, Ohashi T, Nishida K, Nosaki S, Miyakawa T, Tanokura M (2024) BPG4 regulates chloroplast development and homeostasis by suppressing GLK transcription factors and involving light and brassinosteroid signaling. Nature Communications 15: 370

Takagi D, Amako K, Hashiguchi M, Fukaki H, Ishizaki K, Goh T, Fukao Y, Sano R, Kurata T, Demura T, et al (2017a) Chloroplastic ATP synthase builds up a proton motive force preventing production of reactive oxygen species in photosystem I. The Plant Journal 91: 306–324

Takagi D, Ifuku K, Yamori W, Kono M, Ushijima T, Makino A. 2022. Light-harvesting activity is targeted by photosystem I photoinhibition in rice plants irrespective of photosystem I photoinhibitory treatments. Authorea Preprints, DOI: 10.22541/au.166935692.25627459/v1

Takagi D, Ihara H, Takumi S, Miyake C (2019) Growth light environment changes the sensitivity of photosystem I photoinhibition depending on common wheat cultivars. Frontiers in Plant Science 10: 686

Takagi D, Ishizaki K, Hanawa H, Mabuchi T, Shimakawa G, Yamamoto H, Miyake C (2017b) Diversity of strategies for escaping reactive oxygen species production within photosystem I among land plants: P700 oxidation system is prerequisite for alleviating photoinhibition in photosystem I. Physiologia Plantarum 161: 56–74

Takagi D, Tani S (2023) Impact of growth light environment on oxygen sensitivity in rice: Pseudo-first-order response of photosystem I photoinhibition to O_2_ partial pressure. Physiologia Plantarum 175: e14009

Takahashi M, Asada K (1988) Superoxide production in aprotic interior of chloroplast thylakoids. Archives of Biochemistry and Biophysics 267: 714–722

Takeuchi K, Che Y, Nakano T, Miyake C, Ifuku K (2022) The ability of P700 oxidation in photosystem I reflects chilling stress tolerance in cucumber. Journal of Plant Research 135: 681–692

Takeuchi K, Ochiai K, Kobayashi M, Kuroda K, Ifuku K (2024) Light-chilling stress causes hyper- accumulation of iron in shoot, exacerbating leaf oxidative damage in cucumber. Plant and Cell Physiology, 65: 1873–1887

Terashima I, Funayama S, Sonoike K (1994) The site of photoinhibition in leaves of *Cucumis sativus* L. at low temperatures is photosystem I, not photosystem II. Planta 193: 300–306

Terashima I, Kashino Y, Katoh S (1991a) Exposure of leaves of *Cucumis sativus* L. to low temperatures in the light causes uncoupling of thylakoids I. Studies with isolated thylakoids. Plant and Cell Physiology 32: 1267–1274

Terashima I, Matsuo M, Suzuki Y, Yamori W, Kono M (2021) Photosystem I in low light-grown leaves of *Alocasia odora*, a shade-tolerant plant, is resistant to fluctuating light-induced photoinhibition. Photosynthesis Research 149: 69–82

Terashima I, Sonoike K, Kawazu T, Katoh S (1991b) Exposure of leaves of *Cucumis sativus* L. to low temperatures in the light causes uncoupling of thylakoids II. Non-destructive measurements with intact leaves. Plant and Cell Physiology 32: 1275–1283

Tikkanen M, Mekala NR, Aro E-M (2014) Photosystem II photoinhibition-repair cycle protects Photosystem I from irreversible damage. Biochimica et Biophysica Acta (BBA)-Bioenergetics 1837: 210–215

Tiwari A, Mamedov F, Fitzpatrick D, Gunell S, Tikkanen M, Aro E-M (2024) Differential FeS cluster photodamage plays a critical role in regulating excess electron flow through photosystem I. Nature Plants 10: 1592–1603

Trissl H-W (1997) Determination of the quenching efficiency of the oxidized primary donor of Photosystem I, P700^+^: Implications for the trapping mechanism. Photosynthesis Research 54: 237– 240

Tsuyama M, Kobayashi Y (2009) Reduction of the primary donor P700 of photosystem I during steady-state photosynthesis under low light in *Arabidopsis*. Photosynthesis Research 99: 37–47

Tyystjärvi E (2008) Photoinhibition of photosystem II and photodamage of the oxygen evolving manganese cluster. Coordination Chemistry Reviews 252: 361–376

Vass I (2011) Role of charge recombination processes in photodamage and photoprotection of the photosystem II complex. Physiologia Plantarum 142: 6–16

Wada S, Suzuki Y, Miyake C (2020) Photorespiration Enhances Acidification of the Thylakoid Lumen, Reduces the Plastoquinone Pool, and Contributes to the Oxidation of P700 at a Lower Partial Pressure of CO_2_ in Wheat Leaves. Plants 9: 319

Wada S, Yamamoto H, Suzuki Y, Yamori W, Shikanai T, Makino A (2018) Flavodiiron protein substitutes for cyclic electron flow without competing CO_2_ assimilation in rice. Plant Physiology 176: 1509–1518

Wang F, Yan J, Ahammed GJ, Wang X, Bu X, Xiang H, Li Y, Lu J, Liu Y, Qi H (2020) PGR5/PGRL1 and NDH mediate far-red light-induced photoprotection in response to chilling stress in tomato. Frontiers in Plant Science 11: 669

Wang P, Liu W, Han C, Wang S, Bai M, Song C (2024) Reactive oxygen species: multidimensional regulators of plant adaptation to abiotic stress and development. Journal of Integrative Plant Biology 66: 330–367

Yamada Y, Suzuki K, Yanagishita H, Noguchi K (2024) Roles of mitochondrial alternative oxidase in photosynthetic electron transport in illuminated leaves of *Arabidopsis thaliana* at low temperature. Journal of Biosciences 49: 1–12

Yamamoto H, Takahashi S, Badger MR, Shikanai T (2016) Artificial remodelling of alternative electron flow by flavodiiron proteins in *Arabidopsis*. Nature Plants 2: 1–7

Yang Q-Y, Wang X-Q, Yang Y-J, Huang W (2024) Fluctuating light induces a significant photoinhibition of photosystem I in maize. Plant Physiology and Biochemistry 108426

Zhang Y-J, Zhang X, Chen C-J, Zhou M-G, Wang H-C (2010) Effects of fungicides JS399-19, azoxystrobin, tebuconazloe, and carbendazim on the physiological and biochemical indices and grain yield of winter wheat. Pesticide Biochemistry and Physiology 98: 151–157

Zhang Z-S, Jin L-Q, Li Y-T, Tikkanen M, Li Q-M, Ai X-Z, Gao H-Y (2016) Ultraviolet-B radiation (UV-B) relieves chilling-light-induced PSI photoinhibition and accelerates the recovery of CO_2_ assimilation in Cucumber (*Cucumis sativus* L.) leaves. Scientific Reports 6: 34455

Zhang Z-S, Yang C, Gao H-Y, Zhang L-T, Fan X-L, Liu M-J (2014) The higher sensitivity of PSI to ROS results in lower chilling–light tolerance of photosystems in young leaves of cucumber. Journal of Photochemistry and Photobiology B: Biology 137: 127–134

Zivcak M, Brestic M, Kunderlikova K, Sytar O, Allakhverdiev SI (2015) Repetitive light pulse-induced photoinhibition of photosystem I severely affects CO_2_ assimilation and photoprotection in wheat leaves. Photosynthesis Research 126: 449–463

